# 3D Microenvironment-Specific Mechanosensing Regulates Neural Stem Cell Lineage Commitment

**DOI:** 10.1101/2021.09.15.460399

**Authors:** Jieung Baek, Paola A. Lopez, Sangmin Lee, Taek-Soo Kim, Sanjay Kumar, David V. Schaffer

## Abstract

While extracellular matrix (ECM) mechanics strongly regulate stem cell commitment, the field’s mechanistic understanding of this phenomenon largely derives from simplified two-dimensional (2D) culture substrates. Here we found a three-dimensional (3D) matrix-specific mechanoresponsive mechanism for neural stem cell (NSC) differentiation. NSC lineage commitment in 3D is maximally stiffness-sensitive in the range of 0.1-1.2 kPa, a narrower and more brain-mimetic range than we had previously identified in 2D (0.75 – 75 kPa). Transcriptomics revealed stiffness-dependent upregulation of early growth response 1 (*Egr1)* in 3D but not in 2D. *Egr1* knockdown enhanced neurogenesis in stiff ECMs by driving β-catenin nuclear localization and activity in 3D, but not in 2D. Mechanical modeling and experimental studies under osmotic pressure indicate that stiff 3D ECMs are likely to stimulate *Egr1 via* increases in confining stress during cell volumetric growth. To our knowledge, *Egr1* represents the first 3D-specific stem cell mechanoregulatory factor.

---

Mechanical properties of the cellular microenvironment have increasingly been recognized as an important determinant of stem cell behaviors including self-renewal and differentiation^1, 2, 3, 4^. In particular, it has widely been accepted that spatial and temporal variations in extracellular matrix (ECM) stiffness modulate cytoskeletal tension and activate mechanotransductive signaling complexes and transcription factors to regulate stem cell behavior^5, 6, 7, 8^. In our earlier work, we found that the mechanical stiffness of two-dimensional (2D) ECM substrates regulates neural stem cell (NSC) differentiation, where soft ECMs promote neuronal differentiation and stiff ECMs suppress neurogenesis and elevate glial differentiation^9^. These effects were mediated by Rho family GTPase-regulated cellular contractile forces. In addition, the transcriptional coactivator Yes-associated protein (YAP) was upregulated on stiff gels and suppressed neurogenesis by binding and sequestering β-catenin, a transcriptional coactivator that would otherwise upregulate the proneuronal transcription factor NeuroD1^10^.

However, most such in-depth mechanistic analysis of how mechanical cues regulate stem cell behaviors involved 2D platforms, which contrast with natural 3D tissue microenvironments^11, 12^. On 2D matrices, cell spreading and adhesion are polarized and are unlimited by physical confinement, and cells sense ECM stiffness by exerting inward traction forces^13, 14^. In contrast, cell spreading, migration, and growth are confined within 3D matrices, and mechanotransduction can thus also be influenced by other physical factors including the mechanical resistance of the surrounding ECM associated with compression^15, 16, 17^ and degradability^18, 19, 20^. Furthermore, in many 3D contexts, mechanotransduction and force generation occur through focal adhesion-independent mechanisms^20, 21, 22^. For example, cells can migrate in confined 3D matrices in the absence of integrin-mediated adhesions, with cell-ECM forces transmitted through friction^21^. In addition, neural progenitor cell stemness in 3D matrices may be maintained through matrix remodeling-mediated cell-cell contact, in the absence of integrin binding-associated cytoskeletal tension generation^20^. Finally, despite these advances in investigating 3D mechanoregulation, it is not well understood how mechanical inputs ultimately transcriptionally activate target genes that modulate stem cell fate in either 3D or 2D.

In this study, we investigated whether and how NSCs alter their fate choice in response to stiffness in 3D microenvironments. Using engineered hyaluronic acid (HA)-dibenzocyclooctyne (DBCO) hydrogels^23^, we found that soft gels (0.1 kPa) strongly promoted neurogenesis, and stiff gels (1.2 kPa) suppressed neurogenesis, in a stiffness range that is far narrower than we previously found to regulate NSC fate in 2D^9, 10^ and corresponds more closely to the stiffness of adult brain tissue^24, 25, 26^. In addition, RNA-seq revealed that the immediate early gene, early growth response 1 (*Egr1)*^27, 28, 29^, which encodes early growth response protein 1 (EGR1), was highly upregulated on stiff vs. soft gels, in contrast to 2D gels where its expression was negligible. Furthermore, *Egr1* knockdown rescued neurogenesis in stiff gels by reversing its suppression of β-catenin signaling. Finally, a substantial drop in *Egr1* expression with osmotic manipulation of cell volume supports the idea that ECM confining stress during cell volumetric growth in 3D matrices may contribute to 3D stiffness-dependence of *Egr1* expression. In sum, this work implicates *Egr1* as, to our knowledge, the first 3D matrix-specific mechanosensitive regulator of stem cell lineage commitment.

## Results

### NSC lineage commitment in 3D gel is more mechanosensitive than on 2D gels

To investigate whether the lineage commitment of NSCs within 3D matrices is mechanosensitive, we synthesized a series of HA hydrogels in which HA functionalized with DBCO (HA-DBCO) was crosslinked with polyoxyethylene bis(azide), based on strain-promoted alkyne-azide cycloaddition (SPAAC) click chemistry^23^. We engineered these hydrogels to range in stiffness from 0.1-1.2 kPa, which corresponds to the reported elastic modulus of native brain tissue^24, 25, 26^ (**Fig. 1a** and **Supplementary Fig. S1a-f, S2a,b**). To assess the role of integrin ligation, we generated gels that either lacked or included pendant, azide-conjugated RGD peptides (K(N3)GSGRGDSPG), hereafter referred to as RGD- and RGD+ gels, respectively. RGD conjugation did not significantly alter elastic modulus within our working azide: HA monomer range (0.02 - 0.04).

**Fig. 1.**
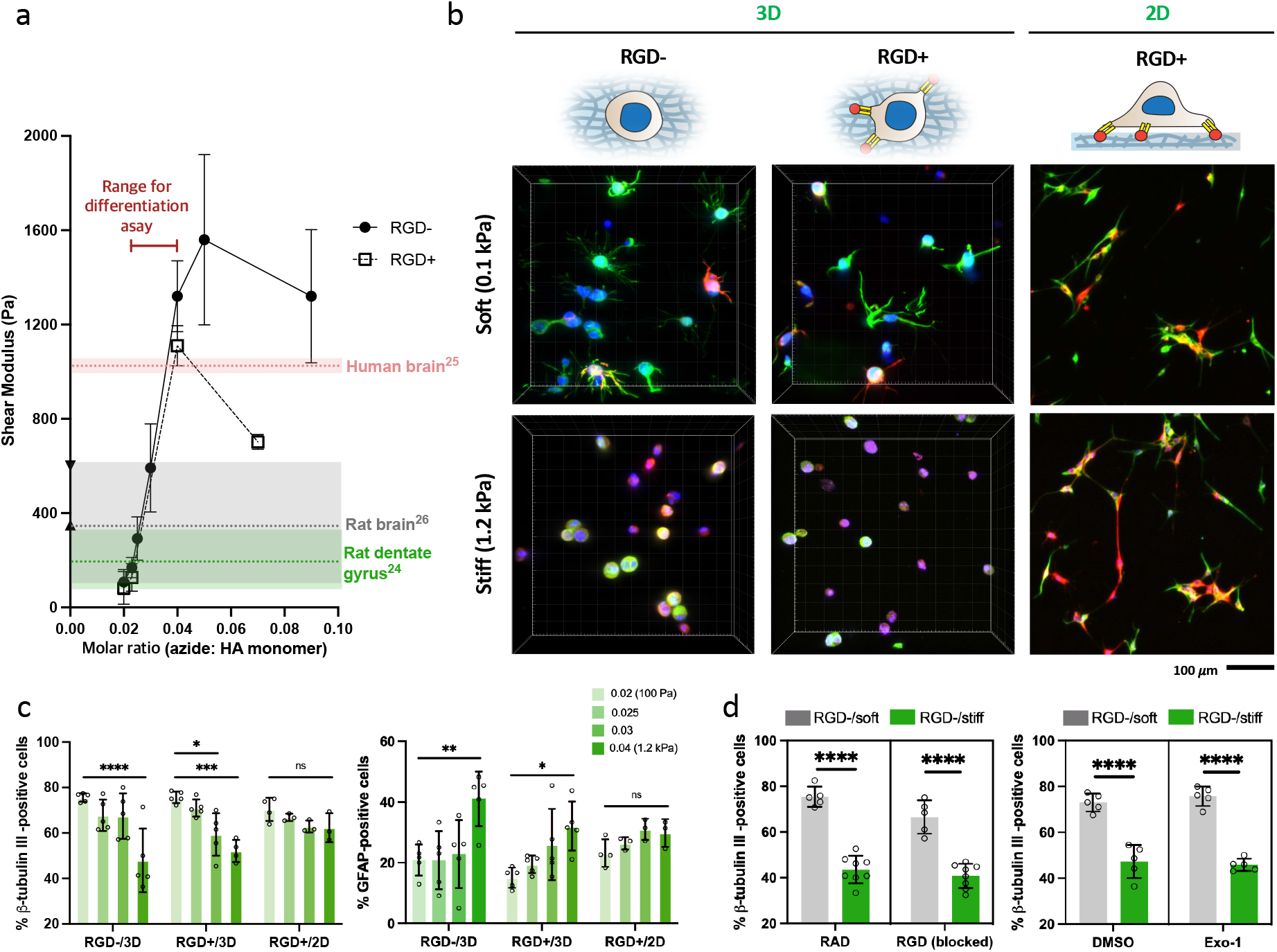
NSC lineage commitment in 3D gels is more mechanosensitive than on 2D gel in 0.1-1.2 kPa. **a**, Shear elastic moduli of hydrogels controlled by the molar ratio of azides (crosslinker) to HA monomers. **b**, Representative images of immunostaining for β-tubulin III (green), GFAP (red), DAPI (blue) in the soft and stiff 3D and 2D hydrogels Scale bar 100 *μ*m. **c**, Quantification of β-tubulin III-positive and GFAP-positive cells in RGD-/3D, RGD+/3D, and RGD+/2D hydrogels. 0.02, 0.025, 0.03, and 0.04 represents the molar ratio of azide (crosslinker) to HA monomer. **d**, Quantification of β-tubulin III-positive cells in the soft (0.1 kPa) and stiff (1.2 kPa) hydrogels with two different condition sets: treatment of RGD sequence-containing peptides and RAD sequence-containing peptides (control) (left) and treatment of Exo-1 and DMSO (control) (right). One-way ANOVA followed by Tukey test \*\*\*\**p*<0.001, \*\**p*<0.01, \**p*<0.05. Graphs show mean ± s.d., n=3 to 5 biological replicates.

We next investigated how NSC lineage commitment was affected by matrix mechanics and RGD status. NSCs were encapsulated and cultured in differentiation medium^9^ that induces a mix of neuronal and glial differentiation for 7 days, fixed, and stained for neuronal (neuron-specific class III β-tubulin, or Tuj1) and astrocytic (glial fibrillary acidic protein, or GFAP) lineage markers. We observed clear stiffness-dependent lineage distribution within this narrow stiffness range, with soft (0.1 kPa) gels strongly promoting neurogenesis and stiff (1.2 kPa) gels suppressing it (**Fig. 1b,c**). As anticipated, the opposite trends were observed with respect to astrocytic differentiation. We also noted that the fraction of cells negative for both Tuj1 and GFAP did not differ significantly within 0.1-1.2 kPa, indicating that this stiffness range does not notably influence overall cell differentiation (**Supplementary Fig. S3**). In addition, active caspase 3 levels for each lineage marker-positive cells were low across all the 3D gel conditions throughout the experiment (<4 %), ruling out the possibility that the observed differences in lineage commitment were due to selective apoptosis of specific lineage progenitors (**Supplementary Fig. S4a-c**). These results differ from our previously reported stiffness-dependent differentiation on 2D gels in two important respects^9^. First, the range of stiffness-sensitivity is much narrower in 3D than in 2D (0.1-1.2kPa for 3D vs. 0.75-75 kPa for 2D), and accordingly when NSCs were cultured on the apical 2D surface of these soft and stiff 3D gel formulations, we found no statistically significant variation in neurogenesis (**Fig. 1c**). Furthermore, only 3D gels showed distinctive morphological differences between soft and stiff gels (**Fig. 1b**), with a higher number of protrusions in soft gels but smaller and more rounded cellular morphologies in stiff gels for both Tuj1+ and GFAP+ cells. Collectively, these results suggest that different mechanistic processes may mediate mechanosensitive lineage commitment in 3D vs. 2D matrices.

Remarkably, very similar stiffness-dependent trends in lineage commitment were observed in both RGD- and RGD+ gels (**Fig. 1c**), implying the dispensability of RGD-integrin ligation to the overall effect. To assess the possibility that cells in RGD-gels may be secreting and engaging RGD-containing proteins, we repeated studies in the presence of soluble blocking RGD peptides (and control RAD peptides), which did not appreciably alter the overall result (**Fig. 1d** and **Supplementary Fig. S5a**). The results were similarly unaffected by treatment with Exo-1 (**Supplementary Fig. S5b)**, which inhibits secretion of proteins including ECM by limiting vesicular trafficking between the ER and Golgi^30^ (**Fig. 1d**). Taken together, these results suggest that the stiffness-dependence of NSC lineage commitment is driven by 3D-specific mechanics within the 0.1-1.2 kPa stiffness range, and that the result is independent of RGD ligand binding.

### *Egr1* expression is dependent on 3D gel stiffness and regulated by 3D matrix-specific mechanics

To investigate molecular mechanisms underlying mechanosensitive lineage commitment in 3D, we performed unbiased RNA sequencing on NSCs encapsulated within the 3D gels under mixed differentiation conditions (**Fig. 2a**). Importantly, we harvested mRNA after 12 hr of following encapsulation, a time that we previously demonstrated NSCs are maximally primed to respond to stiffness cues in 2D^10^. We carried out two comparisons (RGD+/stiff vs. RGD+/soft and RGD-/stiff vs. RGD-/soft) to identify differentially expressed genes (DEGs) between soft and stiff matrices in the presence or absence of RGD. Notably, the heatmap of DEGs for RGD+/stiff vs. RGD+/soft showed similar trends as RGD-gels, i.e. the primary clustering was based on stiffness rather than RGD functionalization (**Fig. 2b**), reenforcing our earlier data (**Fig. 1**) indicating that RGD functionalization is largely dispensable for stiffness-dependent differentiation. Likewise, 83.8 % of DEGs from the comparison between RGD-/stiff vs. RGD-/soft overlapped with those from the RGD+ gels comparison, and the number of DEGs from RGD+ vs. RGD-was negligible for both soft and stiff gels as compared to that for stiff vs. soft. Together, these results indicate that stiffness rather than RGD binding is the more dominant factor regulating the NSC transcriptome at 12 hr in 3D microenvironment.

**Fig. 2.**
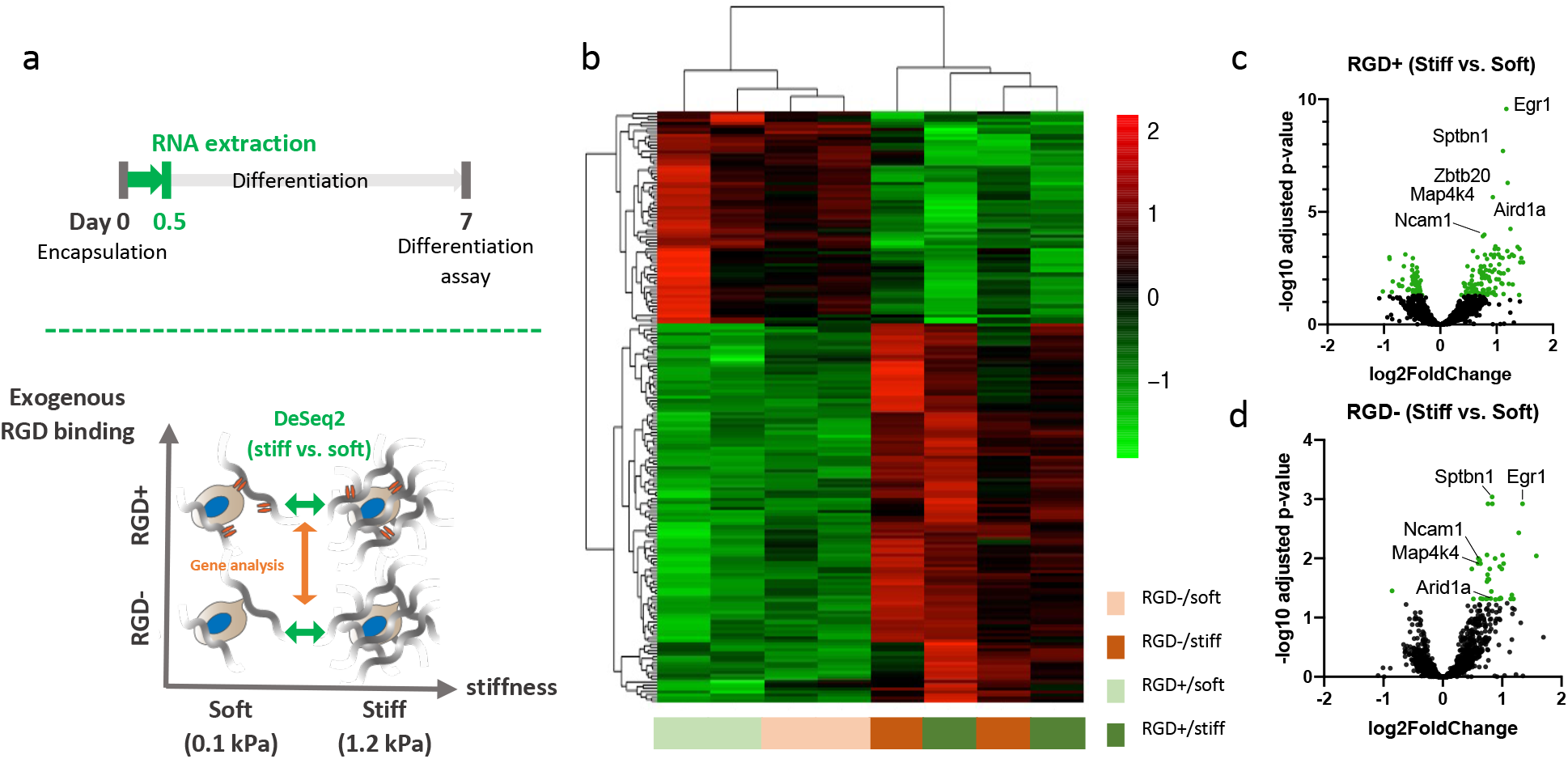
*Egr1* is one of the most differentially expressed genes regulated by the stiffness of 3D matrices. **a**, Schematics of experimental timeline (top) and the strategy to compare the overall transcriptome of NSCs based on the exogenous RGD ligand binding and stiffness (bottom). RNAs isolated from the cells embedded in four different hydrogels (RGD-/soft, RGD-/stiff, RGD+/soft, and RGD+/stiff) were used for this analysis. **b**, A heatmap of the DEGs (from RGD+/stiff vs. RGD+/soft comparison) between the four different hydrogels. Dendrograms indicate the clustering of 3D hydrogel conditions (top) and genes (left). Volcano plots of DEGs from stiff vs. soft hydrogels with (**c**) and without (**d**) RGD functionalization.

Volcano plots of DEGs revealed that compared to soft matrices, stiff matrices uniformly induced higher expression of genes including *Egr1*, Spectrin beta, non-erythrocytic 1 (*Sptbn1*), Mitogen-activated protein kinase kinase kinase kinase-4 (*Map4k4*), Neural Cell Adhesion Molecule 1 (*Ncam1*), and AT-rich interaction domain 1A (*Arid1a*) (**Fig. 2c,d**). The difference was clearest after 12 hours of encapsulation but became more muted after 48 hours, consistent with the idea that NSC lineage is maximally mechanosensitive during a finite time window^10^ (**Supplementary Fig. S6**).

Interestingly, we found that *Egr1* was the gene most differentially expressed between stiff and soft matrices, with the either highest (RGD+) or second highest (RGD-) -log 10 adjusted *p*-value. *Egr1* is an immediate early gene (IEG) tightly associated with neuronal activity as well as a variety of higher order processes such as learning, memory, response to emotional stress, and reward within the central nervous system ^29, 31, 32^. In addition, *Egr1* expression has been reported to be rapidly upregulated by mechanical stimulation in Chinese hamster ovary (CHO) cells^28^ and is a target of the RhoA^33^ and extracellular signal-regulated kinase (ERK)^34^ signaling pathways previously implicated in mechanotransduction. We therefore hypothesized that *Egr1* may be a novel regulator of NSC mechanosensitive lineage commitment in 3D.

Quantitative reverse transcriptase polymerase chain reaction (qRT-PCR) validation of the RNA-seq results demonstrated that stiffness-dependent *Egr1* expression appeared within 5 hr of encapsulation at levels two-fold higher in stiff as compared to soft gels (**Fig. 3a**). In addition, this level increased for all 3D gels with encapsulation times up to 12 hr. Notably, *Egr1* levels were 70 and 1500 times higher in 3D gels than on corresponding 2D gels cultured in otherwise identical conditions after 5 hr and 12 hr, respectively, and *Egr1* upregulation was evident only in 3D. A similar trend was observed irrespective of material platform or RGD incorporation, with negligible *Egr1* expression on 2D gels even when stiffness was raised to 73 kPa (**Supplementary Fig. S7a**). As with lineage commitment, addition of soluble RGD peptides or Exo-1 did not significantly affect the 3D stiffness dependence of *Egr1* (**Fig. 3b,c**). To confirm stiffness-dependent expression of *Egr1* at the protein level, we performed western blotting for EGR1 in the four hydrogels (RGD-/soft, RGD-/stiff, RGD+/soft, RGD+/stiff). We observed the same trends as in RNA-seq and qRT-PCR (**Fig. 2c,d** and **Fig. 3a**), with markedly higher expression level in stiff gels (**Fig. 3d** and **Supplementary Fig. S7b,c**). EGR1 protein expression was also negligible in 2D gels, regardless of material platform and RGD incorporation even at higher stiffness (73 kPa) (**Supplementary Fig. S7d,e**). To summarize, *Egr1* expression depended on stiffness only in 3D matrices and independently of RGD binding (**Fig. 3e**), with a trend that corresponded well with cell differentiation (**Fig. 1b-d**).

**Fig. 3.**
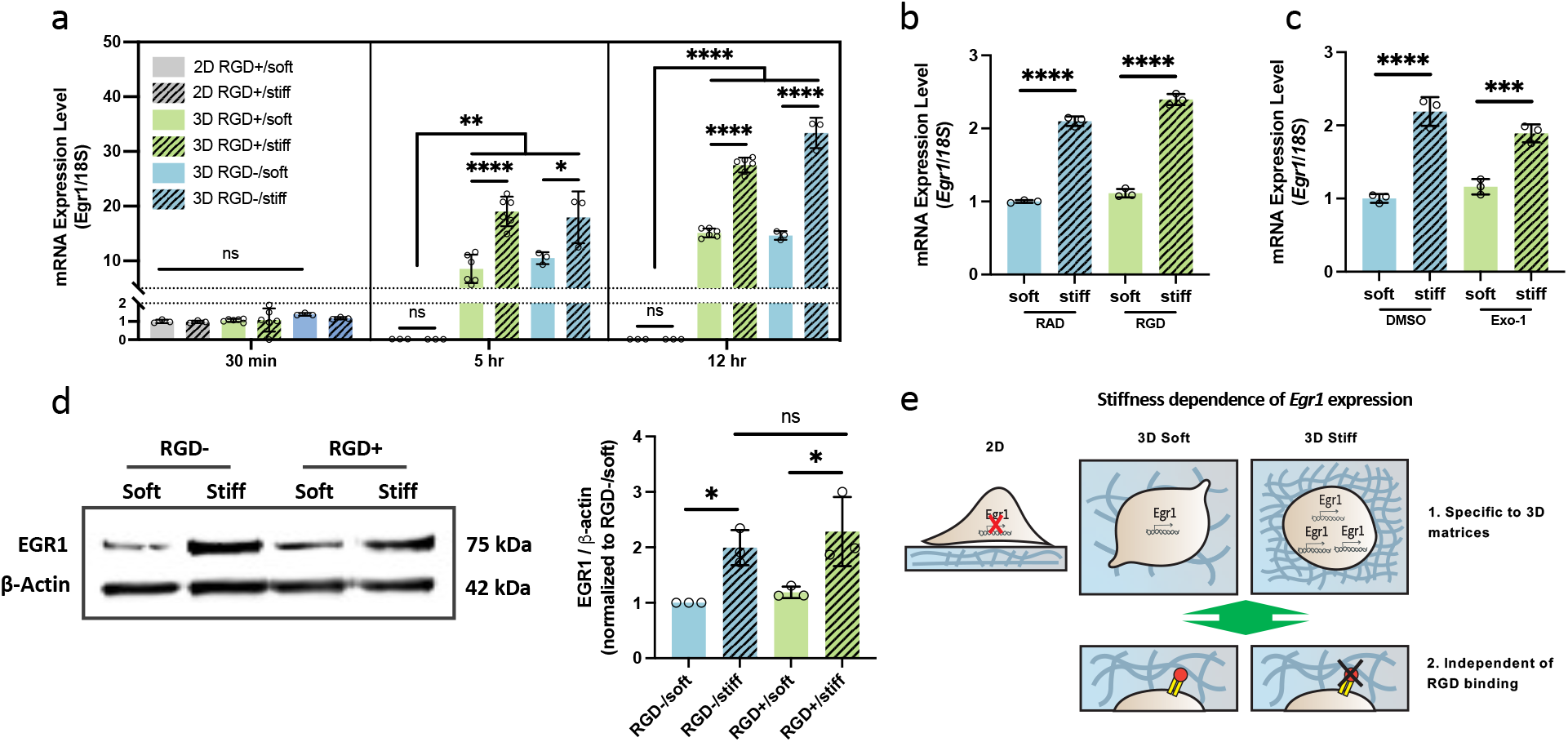
*Egr1* mRNA expression is regulated by 3D matrices-specific mechanics. **a**, *Egr1* mRNA expression kinetics during differentiation within bare (RGD-) hydrogels and RGD-ligated (RGD+) 2D and 3D hydrogels. Each level is relative to the expression level on 2D soft gel right after encapsulation (30 min). Hydrogels of 0.1 kPa and 1.2 kPa were used for soft and stiff condition, respectively. Expression of mRNA level for *Egr1* in the bare soft and stiff hydrogels with RGD sequence-containing peptides or RAD-containing peptides (control) (**b**) and with DMSO (control or Exo-1 (**c**). **d**, Western blot and quantification of EGR1 protein expression in NSCs encapsulated within the four different hydrogels (RGD-/soft, RGD-/stiff, RGD+/soft, and RGD+/stiff) for 24 hr. **e**, Schematic illustration showing the characteristics of stiffness-dependence of *Egr1* transcription in 3D gels. The dependence was observed only in 3D matrices independently of RGD-integrin binding. One-way ANOVA followed by Tukey test \*\*\*\**p*<0.001, \*\*\**p*<0.005, \**p*<0.05. Graphs show mean ± s.d.

### EGR1 plays a role in the stiffness dependence of NSC lineage commitment only in 3D matrices by regulating β-catenin signaling

To investigate the potential importance of *Egr1* in stiffness-dependent NSC fate commitment in 3D, we depleted *Egr1* with lentivirally-delivered short hairpin RNAs (shRNAs) (**Fig. 4a**). Two shRNAs (shEGR1-1 and shEGR1-2) targeting different regions of *Egr1* mRNA efficiently knocked down EGR1 protein expression compared to cells transduced with a control shRNA (shCtrl) (**Fig. 4b**). These cells were then differentiated within four different 3D gels (RGD-/soft, RGD-/stiff, RGD+/soft, and RGD+/stiff) under mixed differentiation conditions. Notably, *Egr1* knockdown rescued neurogenesis in cells in stiff gels compared with shCtrl and naïve cells (**Fig. 4c,d**) to levels similar to what we observed for naïve and shCtrl cells in soft matrices, again independent of RGD. Interestingly, no significant difference in neurogenesis among naïve, shCtrl, and shEGR1-1 cells was detected for cells on 2D gels (**Fig. 4e,f**). This finding is consistent with the very low, stiffness-independent *Egr1* expression seen earlier on 2D gels and further reinforces that *Egr1* does not mediate mechanosensitive lineage commitment on 2D gels (**Fig. 4g**).

**Fig. 4.**
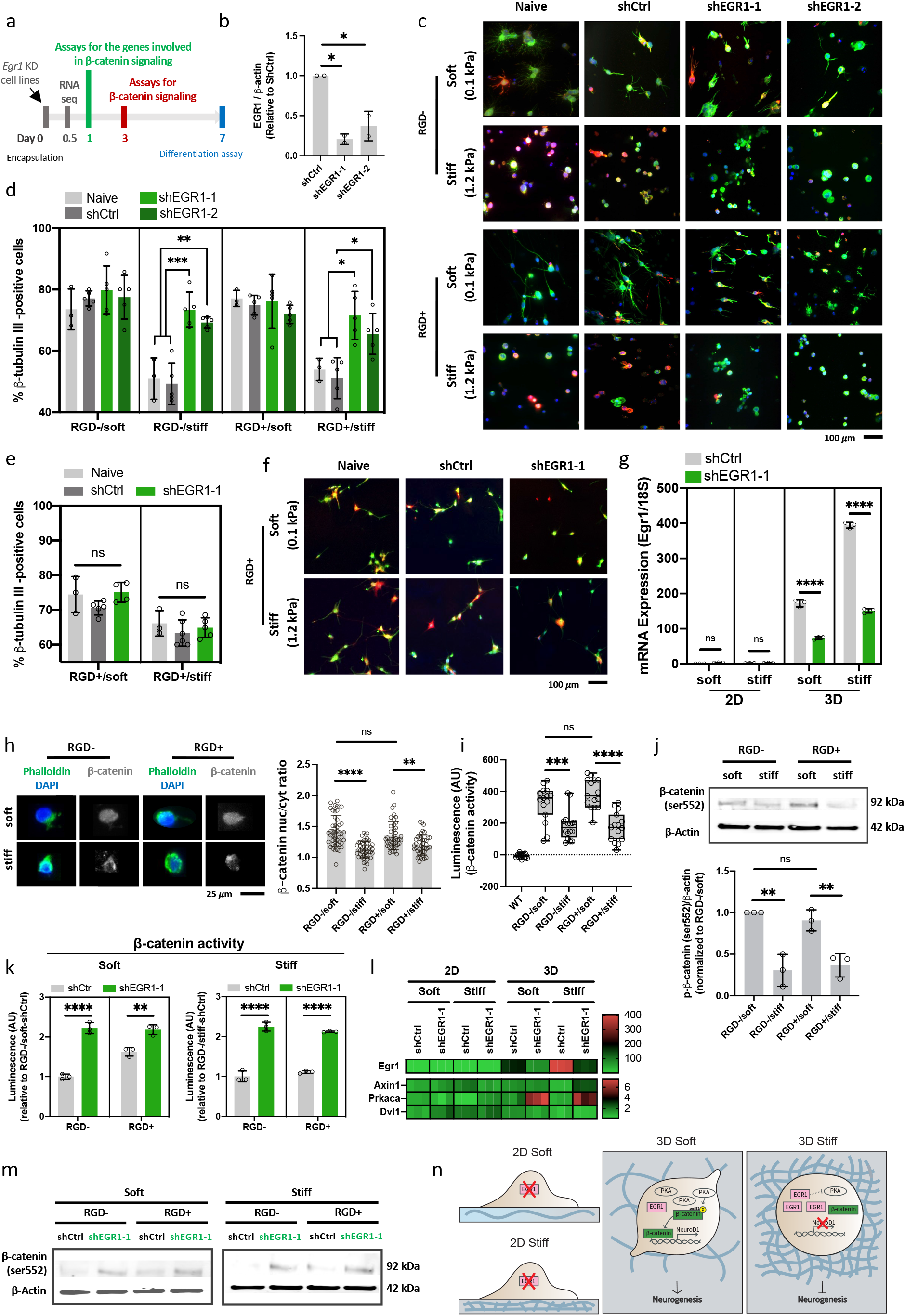
EGR1 plays a role in mechanosensitive NSC lineage commitment only in 3D matrices by regulating β-catenin signaling. **a**, Experimental timeline of revealing the mechanism how EGR1 regulates NSC lineage commitment. **b**, EGR1 protein expression after functional depletion using shRNA. n=2 biological replicates. Representative immunofluorescence images of naïve, shCtrl, and *Egr1* knockdown cell lines stained for β-tubulin III (green), GFAP (red), DAPI (blue) and quantification of the neurogenesis in 3D (**c, d**) and 2D (**e, f**) soft (0.1 kPa) and stiff (1.2 kPa) hydrogels. Scale bar, 100 *μ*m. n=3 to 5 biological replicates. **g**, mRNA expression level of *Egr1* in shCtrl and shEGR1-1 cell lines in 2D and 3D gels. **h**, Representative images (left) of immunofluorescence staining for β-catenin (gray), F-actin (green), nucleus (blue) and quantification (right) of the β-catenin nuclear localization for the dissected NSCs encapsulated with four different hydrogels: RGD-/soft, RGD-/stiff, RGD+/soft, RGD+/stiff. scale bar 50 *μ*m. n>41 cells per group was used for the quantification. **i**, Luciferase assay for β-catenin activity in WT and NSC reporter cells embedded in the four different 3D gels. n=15 technical replicates including n=3 biological replicates per each condition. **j**, Western blotting for the active β-catenin (phosphorylated at Ser552) of the cells embedded in the four different gels. **k**, Luciferase assay showing β-catenin activity of shCtrl and *Egr1* knockdown cell lines in the 3D hydrogels. Each value is relative to that of shCtrl cells in RGD-gels for soft and stiff conditions separately. **l**, mRNA expression level (qRT-PCR) of *Egr1* and three different genes (*Axin1, Prkaca, Dvl1*) involved in Wnt signaling in 2D and 3D gels. All the level for each gene is relative to that of shCtrl on 2D soft gels. n=3 biological replicates. **m**, Western blotting of active β-catenin (Ser552) of shCtrl and shEGR1-1 cells encapsulated with the four different gels. **n**, Schematic of the suggested mechanism how differentially expressed EGR1 in soft and stiff matrices leads to the stiffness-dependent lineage commitment in 3D matrices. One-way ANOVA followed by Tukey test \*\*\*\**p*<0.001, \*\*\**p*<0.005, \*\**p*<0.01, \**p*<0.05. Graphs show mean ± s.d.

We next asked how *Egr1* regulates neurogenesis in 3D. The Wnt/β-catenin signaling pathway is known to play critical roles in development, differentiation, and maintenance of stemness ^35, 36, 37^, and our and other groups have implicated β-catenin signaling in NSC differentiation into neurons^10, 35, 38^. Active, nuclearly localized β-catenin transcriptionally activates *NeuroD1*, a proneuronal transcription factor for NSCs. Intriguingly, it has been reported that EGR1 binding sites are present in the promoters of >15 genes encoding factors in Wnt signaling pathway, indicating a potential link between *Egr1* and Wnt signaling in NSC differentiation^27^.

To investigate whether β-catenin can regulate neurogenesis within 3D matrices, CHIR, a highly potent and specific GSK3 inhibitor that potentiates β-catenin activity, was added for 72 hr under differentiation conditions. CHIR treatment enhanced neuronal differentiation and reduced astrocytic differentiation in stiff gels (**Supplementary Fig. S8a-f**). We then asked if β-catenin nuclear localization is regulated by 3D gel stiffness (**Fig. 4h**) and found that its nuclear/cytoplasmic intensity ratio was higher in both RGD-/soft and RGD+/soft gels compared to the stiff gels, implying that β-catenin more strongly traffics to the nucleus in soft than in stiff gels. We then directly assessed β-catenin-dependent transcription using an established β-catenin-responsive luciferase reporter^39^, 7xTFP, which we stably introduced into NSCs. We validated this reporter in our system by treating NSCs with CHIR, which resulted in a dose-dependent increase in bioluminescence (**Supplementary Fig. S8a**). After 72 hr of differentiation in 3D gels, higher luciferase expression was observed in soft gels than in stiff gels (**Fig. 4i**). Furthermore, western blotting revealed that the soft gels exhibited higher activated β-catenin (phosphorylated at Ser552) than in stiff gels, confirming stiffness-dependent β-catenin activation at the protein level (**Fig. 4j**). Taken together, these observations demonstrate that stiffness-dependent neurogenesis in 3D matrices is regulated by β-catenin signaling. Importantly, the stiffness-dependent β-catenin activity was not accompanied by differences in levels of YAP, a transcriptional co-activator that has previously been implicated in stiffness-dependent stem cell differentiation^40, 41, 42^ (**Supplementary Fig. S8g**). We recently showed that YAP is upregulated in NSCs on 2D stiff gels where it suppresses neurogenesis by binding and sequestering active β-catenin^10^. The absence of significant stiffness-dependent YAP expression levels in 3D further supports a distinct mechanism for mechanosensitive lineage commitment in 2D vs. 3D.

To determine whether *Egr1* functionally regulates β-catenin signaling, we made additional shCtrl and shEGR1-1 cell lines that express the 7xTFP β-catenin-responsive luciferase reporter (**Supplementary Fig. S8h**). *Egr1* suppression greatly enhanced β-catenin activity in all gels, independent of stiffness or RGD functionalization (**Fig. 4k**). Interestingly, even though *Egr1* suppression increased β-catenin activation in both soft and stiff gels, neurogenesis was not enhanced in soft gels (**Fig. 4d**), suggesting that β-catenin signaling in 3D soft gels may already be functionally saturated.

To investigate how EGR1 may affect β-catenin signaling in 3D gels, we performed qPCR to quantify mRNA expression levels of three genes that are involved in Wnt signaling and whose promoters harbor EGR1 binding sites^27^: *Axin1*, Protein kinase cAMP-activated catalytic subunit alpha (*Prkaca*), and Dishevelled segment polarity protein 1 (*Dvl1*) (**Fig. 4l**). Notably, *Egr1* knockdown did not reduce *Axin1* levels despite reports that it upregulates this Wnt signaling repressor and did not significantly alter *Dvl1* levels; however, cells embedded in 3D gels showed 3.9-to 5.5-fold enhancement in the expression of *Prkaca* after *Egr1* knockdown. *Prkaca* encodes protein kinase A (PKA), which has been reported to phosphorylate β-catenin at Ser552 and Ser675 and thereby promote its transcriptional activity^43, 44, 45, 46^. Consistent with this possibility, *Egr1* knockdown increased the level of Ser552-phosphorylated β-catenin for all the four different gel conditions (**Fig. 4m**), suggesting that *Egr1* upregulation in stiff gels suppresses *Prkaca*, thereby decreasing active β-catenin (Ser552). Notably, *Egr1* depletion in 2D did not significantly affect the expression of *Axin1, Prkaca*, or *Dvl1* (**Fig. 4l**), corresponding well with our findings that *Egr1* knockdown influences NSC fate decisions in 3D but not 2D (**Fig. 4d,e**).

Collectively, our results suggest a mechanism by which the stiffness-dependent *Egr1* expression regulates neurogenesis: the abundance of EGR1 in stiff gels may suppress *Prkaca* expression and thus β-catenin signaling to reduce neuronal differentiation in 3D matrices (**Fig. 4n**). Additionally, the vanishingly low levels and lack of stiffness-dependent *Egr1* expression in 2D are consistent with weak mechanoregulation in the 0.1-1.2 kPa stiffness range (**Fig. 1c**).

### Higher *Egr1* expression in 3D stiff gels is associated with cytoskeletal assembly

We next investigated mechanisms that may link 3D matrix stiffness to *Egr1* expression. We and others have strongly implicated cytoskeletal assembly and tension in 2D stiffness-dependent stem cell lineage commitment^2, 9, 47^. Cross-sectional imaging in 3D gels showed that cells had more condensed actin structures in stiff than in soft gels (**Fig. 5a**). Conversely, cells on 2D gels did not exhibit dramatic differences in actin assembly within same stiffness range, though cells on gels stiffer than ∼2 kPa began to show higher spreading with enhanced actin intensity. These results indicate that the stiffness range of 0.1 – 1.2 kPa is sufficient to drive stiffness-dependent changes in actin cytoskeletal changes in 3D but not 2D. Furthermore, quantification of peak cortex actin intensity after linearization (**Fig. 5b**) revealed that cells in stiff gels showed higher cortical actin intensity than those in soft gels, while exhibiting no striking difference in actin thickness **(Supplementary Fig. S9a**). To investigate how cytoskeletal tension regulates *Egr1* expression, cells in 3D gels were treated with small molecule inhibitors against myosin II, actin polymerization, and focal adhesion kinase (FAK) (**Fig. 5c** and **Supplementary Fig. S9b,c**). Interestingly, inhibition of myosin II and FAK by blebbistatin and PF, respectively, did not substantially alter the stiffness dependence of *Egr1* expression. However, inhibition of actin polymerization by cytochalasin D (cyt D), which lowered the level of actin intensity but not the thickness (**Fig. 5d** and **Supplementary Fig. S9d**), decreased *Egr1* expression and weakened its stiffness dependence. Consistently, disruption of actin polymerization rescued lower neurogenesis in stiff gels (**Fig. 5e**). These results suggest that actin assembly may play a role in stiffness-dependent *Egr1*-mediated neurogenesis that is independent of myosin II-mediated contraction and FAK signaling (**Fig. 5f**).

**Fig. 5.**
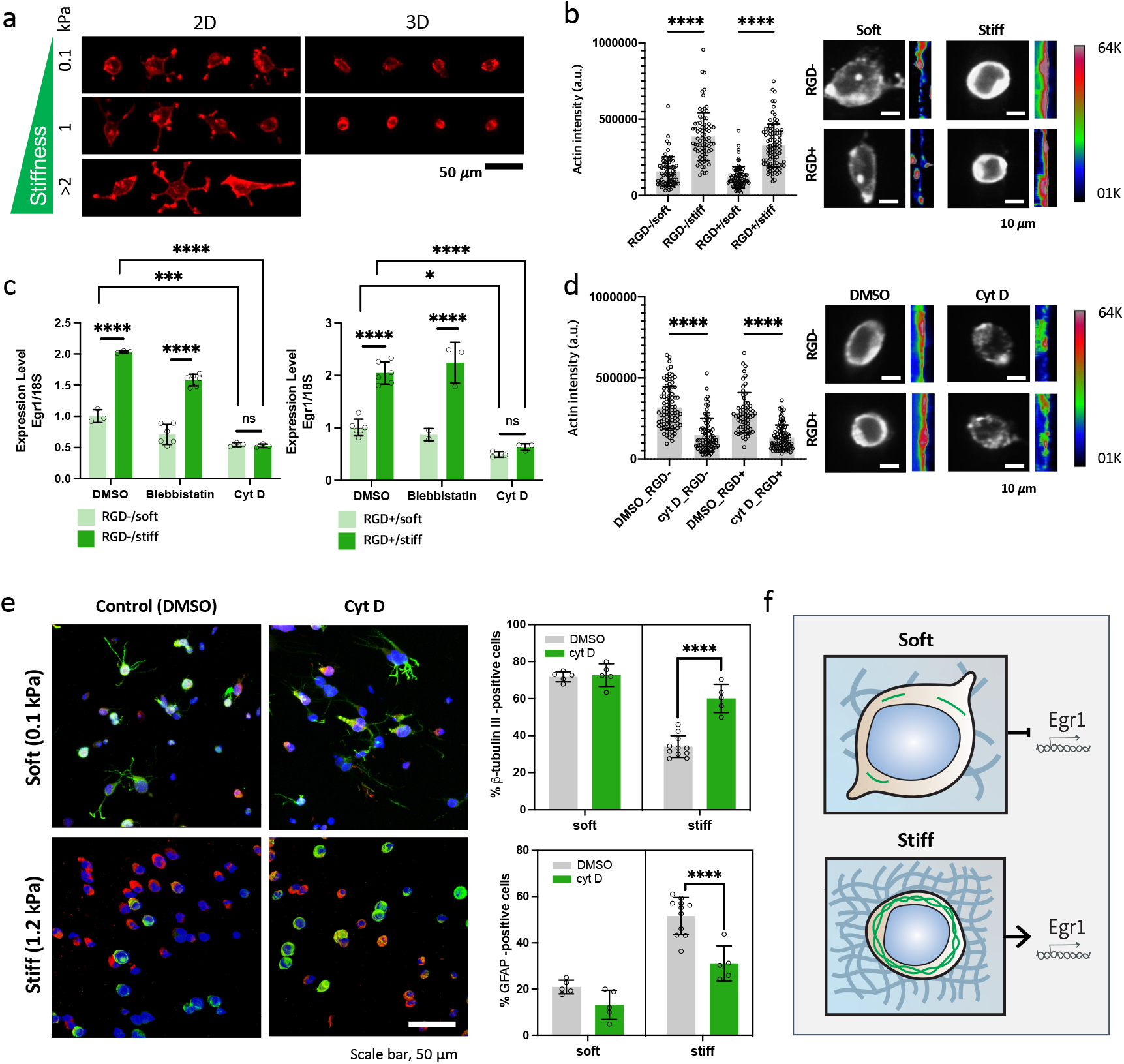
Actin formation regulates stiffness-dependence of *Egr1* expression in 3D matrices. **a**, Representative images of the Rhodamine-phalloidin-stained NSCs differentiated in 2D and 3D gels for 4 hr with different stiffness (0.1 kPa, 1.2 kPa, >2 kPa). All the gels were RGD-functionalized. Images for 3D gels were obtained after dissection. Scale bar 50 *μ*m. **b**, Quantification of peak cortex actin intensity line-scan after background subtraction (left) and representative images of phalloidin-stained cells in the four different 3D gels and color-coded representation of linearized and zoomed-in view of the cortex for each image (right). Images for 3D gels were obtained after dissection. Scale bar 10 *μ*m. **c**, mRNA expression level of *Egr1* in the cells after 5 hr of encapsulation with RGD-(left) and RGD+ (right) gels after treatment of blebbistatin (1 *μ*M) and cytochalasin D (1 *μ*M) (n=3). DMSO was treated as control. **d**, Quantification of peak cortex actin intensity line-scan (left) and images of phalloidin-stained cells in 3D stiff gels under the DMSO-(control) and cytochalasin D-treated conditions (right). **e**, Representative images of immunostaining for β-tubulin III (green), GFAP (red), DAPI (blue) and quantification of β-tubulin III- and GFAP-positive cells in RGD+ 3D hydrogels after treatment of DMSO (control) and cytochalasin D. Scale bar 50 *μ*m. **f**, Schematics showing that actin formation regulates stiffness-dependent *Egr1* expression in 3D matrices. One-way ANOVA followed by Tukey test \*\*\*\**p*<0.001, \*\*\**p*<0.005, \*\**p*<0.01, \**p*<0.05. Graphs show mean ± s.d.

### Confining stress is a 3D gel-specific mechanism that may contribute to stiffness-dependent actin assembly and *Egr1* expression

Given the 3D-specific role of *Egr1*, we next considered biophysical mechanisms of mechanosensing that may be particularly important in 3D matrices. We reasoned that confining stresses experienced by cells as their volumes expand against the mechanical resistance of the matrix ^15, 48^ could represent an important regulatory factor operant in 3D but not 2D (**Fig. 6a**). We observed that cell volume increases with encapsulation time following induction of differentiation for all the four 3D gels: RGD-/soft, RGD-/stiff, RGD+/soft, and RGD+/stiff (**Fig. 6b**). The soft gels showed slightly higher initial volume and more rapid volumetric growth than stiff gels for both RGD- and RGD+ conditions. This trend was also observed for both soluble RAD peptide- and RGD peptide-treated conditions (**Supplementary Fig. S10**), demonstrating that volumetric growth within 3D gels occurs in both soft and stiff gels and is independent of RGD-integrin binding.

**Fig. 6.**
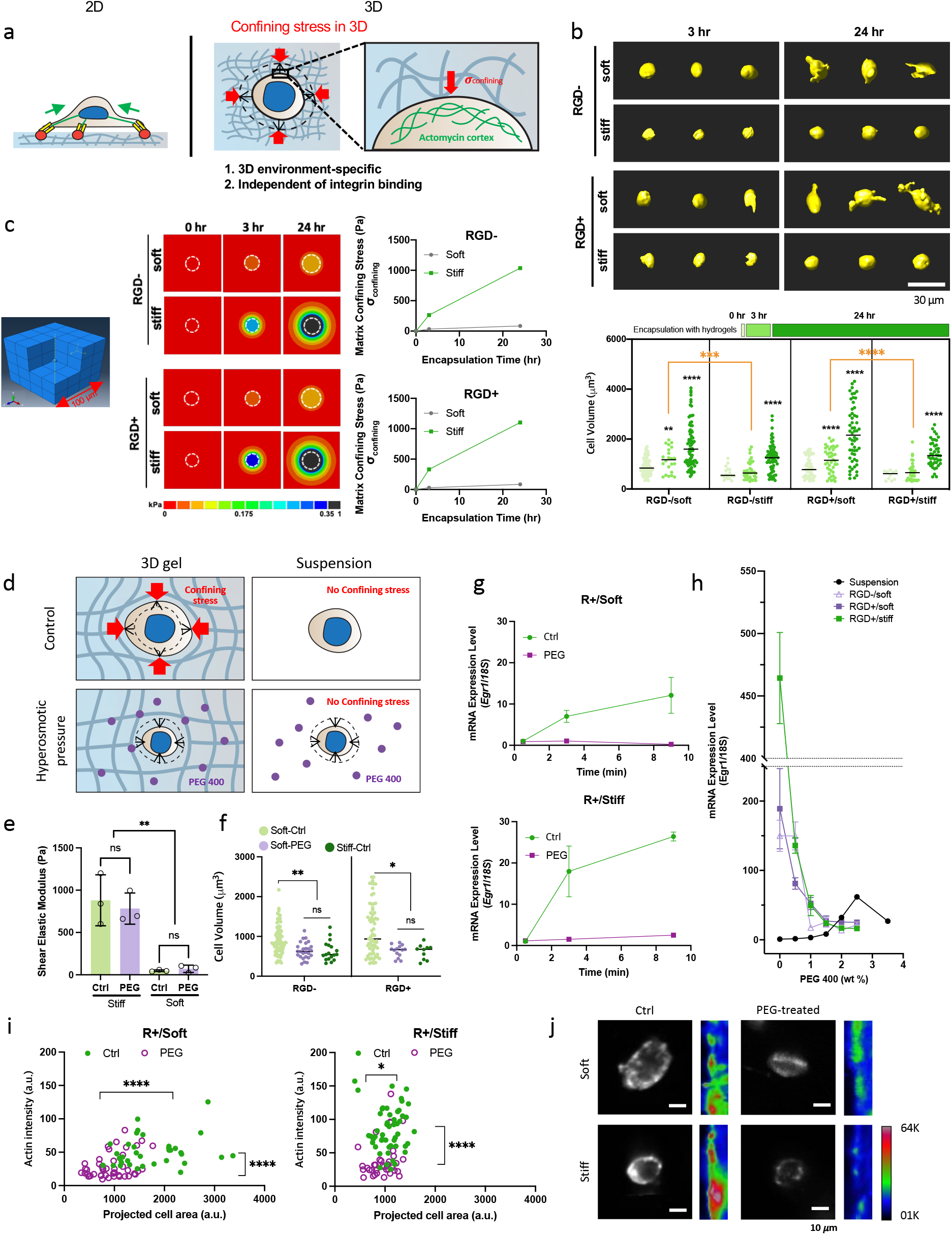
Confining stress during cell volumetric may play an important role in stiffness-dependent *Egr1* expression. **a**, Schematics suggesting confining stress as a 3D gel-specific and RGD binding-independent physical factor that may regulate *Egr1* expression in 3D gels. **b**, Representative 3D rendering of single NSCs stained with cell membrane dye (R18) after 3 hr, 24 hr of 3D gel encapsulation (top) and quantification of cell volumes (bottom). The cells were embedded in the four different 3D gels (RGD-/soft, RGD-/stiff, RGD+/soft, RGD+/stiff) under the spontaneous differentiation condition. 10-77 cells were used for the quantification. Scale bar 30 *μ*m. **c**, ABAQUS simulation for calculating the confining stress of matrix exerted to the cells during cell volumetric growth in the four different 3D gels. Defined model system for the simulation (left), color-coded stress field with the direction of matrix to the cells (center), and quantified stress values with incubation time (right). White dot line represents the cell boundary. **d**, Schematics of applying hyperosmotic pressure to the cells in 3D gel to release the effect of confining stress during cell volumetric growth. Dotted line represents the cell size right after encapsulation. **e**, Shear elastic moduli of RGD+/soft and RGD+/stiff gels incubated under non-treated (Ctrl) and PEG 400 (1.5 wt%)-treated conditions for 3 hr. **f**, Quantification of cell volume in 3D gels for 3 hr under three different conditions: soft gels with and without PEG 400 (1.5 wt%) treatment and stiff gels without PEG 400. **g**, Change of *Egr1* mRNA expression level during differentiation within soft and stiff RGD+ 3D hydrogels under non-treated (Ctrl) and PEG 400 (1.5 wt%)-treated conditions (30 min, 3 hr, and 9 hr). **h**, *Egr1* mRNA expression level with increasing PEG concentration for both cell suspension and the cells in 3D gels. **i**, Scatter plot of peak cortex actin intensity versus projected cell area of the cells encapsulated in 3D RGD+/soft (left) and RGD+/stiff (right) gels under non-treated (Ctrl) or PEG 400 (1.5 wt%)-treated condition for 3 hr. **j**, Representative images of phalloidin-stained cells in 3D RGD+ gels under the Ctrl and PEG-treated conditions and color-coded representation of linearized and zoomed-in view of the cortex for each image (right). Images for 3D gels were obtained after dissection. Scale bar 10 *μ*m. Statistical significance was determined by Student’s t-test (two-tail) between two groups (**i**), and three or more groups were analyzed by One-way ANOVA followed by Tukey test \*\*\*\**p*<0.001, \*\*\**p*<0.005, \*\**p*<0.01, \**p*<0.05. Graphs show mean ± s.d.

We next calculated the confining stress exerted from the surrounding gels to the cell during cell volumetric growth by finite element modeling using ABAQUS 6.14 (**Fig. 6c**). We defined a model system with a thermally expanding sphere embedded with an elastic cube, where the spheres represent cells that volumetrically grow at a morphological aspect ratio of 1 and a zero stress just after encapsulation (0 hr). Intriguingly, stiff gels showed approximately 8- and 11-times greater stress than soft gels after only 3 hr of encapsulation under RGD- and RGD+ conditions, respectively. Furthermore, the time course of increasing stress with encapsulation time corresponds with our observed kinetics for *Egr1* upregulation (**Fig. 3a**).

We next investigated whether *Egr1* expression is altered after relieving the effect of ECM confining stress (**Fig. 6c,d**). Cell volumetric growth was inhibited by applying an osmotic stress by adding 400-Da polyethylene glycol (PEG 400) to the culture medium. To account for potential ECM-independent effects of osmotic pressure on cells, suspension cells were also incubated under osmotic pressure. Rheometric analysis of the gel under non-treated (Ctrl) and PEG 400-treated (PEG) conditions confirmed that 1.5 wt% of PEG does not significantly affect the shear elastic moduli of both soft (0.1 kPa) and stiff (1 kPa) gels (**Fig. 6e**). However, PEG treatment limited the cell volumetric expansion within the soft gels, even resulting in the volume similar to those in stiff gels for both RGD- and RGD+ conditions after 3 hr (**Fig. 6f**). Interestingly, the restricted cell volume expansion limited the increase of *Egr1* expression level with 3D gel encapsulation time compared to the unrestricted (Ctrl) condition, ultimately leading to significantly different *Egr1* levels between the Ctrl and osmotic pressure (PEG) conditions after 9 hr (**Fig. 6g**). This volumetric restriction resulted in a remarkable 8-18 fold drop in *Egr1* expression level in a PEG concentration-dependent manner after 3 hr of encapsulation, whereas suspension cells in contrast showed slight increase in *Egr1* expression with PEG 400 concentration (**Fig. 6h**). This slight *Egr1* increase indicates that a decrease in cell volume, under suspension conditions where cells do not interact with ECM, actually enhances *Egr1* expression. Similar results were observed with 2D gels, where PEG treatment induced a 10-20 fold *Egr1* increase **(Supplementary Fig. S11a,b**). These results are consistent with the hypothesis that ECM confining stress during cell volumetric growth, a 3D matrix-specific physical factor independent of volumetric growth alone, may play an important role in *Egr1* expression in 3D microenvironments.

Since we previously observed that actin assembly influences *Egr1* expression (**Fig. 5c**), we next examined if there is a change in actin architecture after manipulating osmotic pressure. Cells in both soft and stiff 3D gels showed an overall reduction of cortex actin intensity under osmotic pressure (**Fig. 6i,j**). This trend correlates well with the lower *Egr1* expression level seen under osmotic pressure (**Fig. 6g**), suggesting a possible causal link between constrained volumetric growth-mediated regulation of *Egr1* and cytoskeletal assembly.

### Stiffness-dependent *Egr1* expression is associated with H3K9 trimethylation

In addition to cytoskeletal reorganization, cells could also potentially respond to confining stress through nuclear reorganization. Stiff gels induced a smaller nuclear size than soft gels for both RGD- and RGD+ conditions, indicating that nucleus is mechanically influenced by 3D gel stiffness regardless of RGD binding, as we previously observed for actin formation (**Fig. 7a**). Chromatin architecture, which is regulated in part by enzymatic acetylation and methylation, has recently been reported to exhibit stiffness-dependent accessibility in 3D matrices, with concomitant changes in gene expression^49^. In particular, demethylation of histone H3 lysine-9 trimethylation (H3K9me3) has been shown to induce Pol II recruitment and increase *Egr1* transcription in cells under force^28^. It has recently been reported from Hi-C analysis that H3K9me3 is a strong functional marker for transcriptionally inactive chromosomal regions^50^. Accordingly, we examined whether H3K9 trimethylation levels are dependent on stiffness under three different gel conditions: RGD-presenting 2D gels, RGD- and RGD+ 3D gels (**Fig. 7b**). Intriguingly, 3D but not 2D gels exhibited stiffness-dependent H3K9 methylation, with higher H3K9me3 in soft than in stiff gels. Furthermore, the H3K9 demethylase inhibitor JIB-04 strikingly reduced *Egr1* expression for all 3D gel conditions (RGD-/soft, RGD-/stiff, RGD+/soft, and RGD+/stiff), despite an only 1.3-fold increase in H3K9me3 levels after JIB-04 treatment (**Fig. 7c** and **Supplementary Fig. S12**). Stiffness-dependent *Egr1* expression was also slightly weakened by inhibition of H3K9 demethylase. Although this value was not as significantly altered as with actin assembly inhibition (**Fig. 5c**), the substantial decrease in *Egr1* expression level for each gel condition indicates that H3K9me3 restricts the transcription of *Egr1* in 3D gels. Given that both actin and H3K9 trimethylation play a role in *Egr1* expression in 3D gels, we investigated whether altered actin assembly and H3K9 trimethylation are directly correlated with each other in a process of regulating stiffness-dependent *Egr1* transcription in 3D gels. H3K9me3 levels did not substantially change after inhibition of actin assembly in either RGD- or RGD+ stiff gels, indicating that any alterations in cortical actin assembly observed in 3D stiff gels do not significantly influence H3K9 trimethylation (**Fig. 7d**). Taken together, our results demonstrate that less trimethylation of H3K9 in stiffer gels may induce higher *Egr1* expression levels compared to soft gels, an effect that could potentially be linked with the stiffness-dependent confining stress (**Fig. 7e**).

**Fig. 7.**
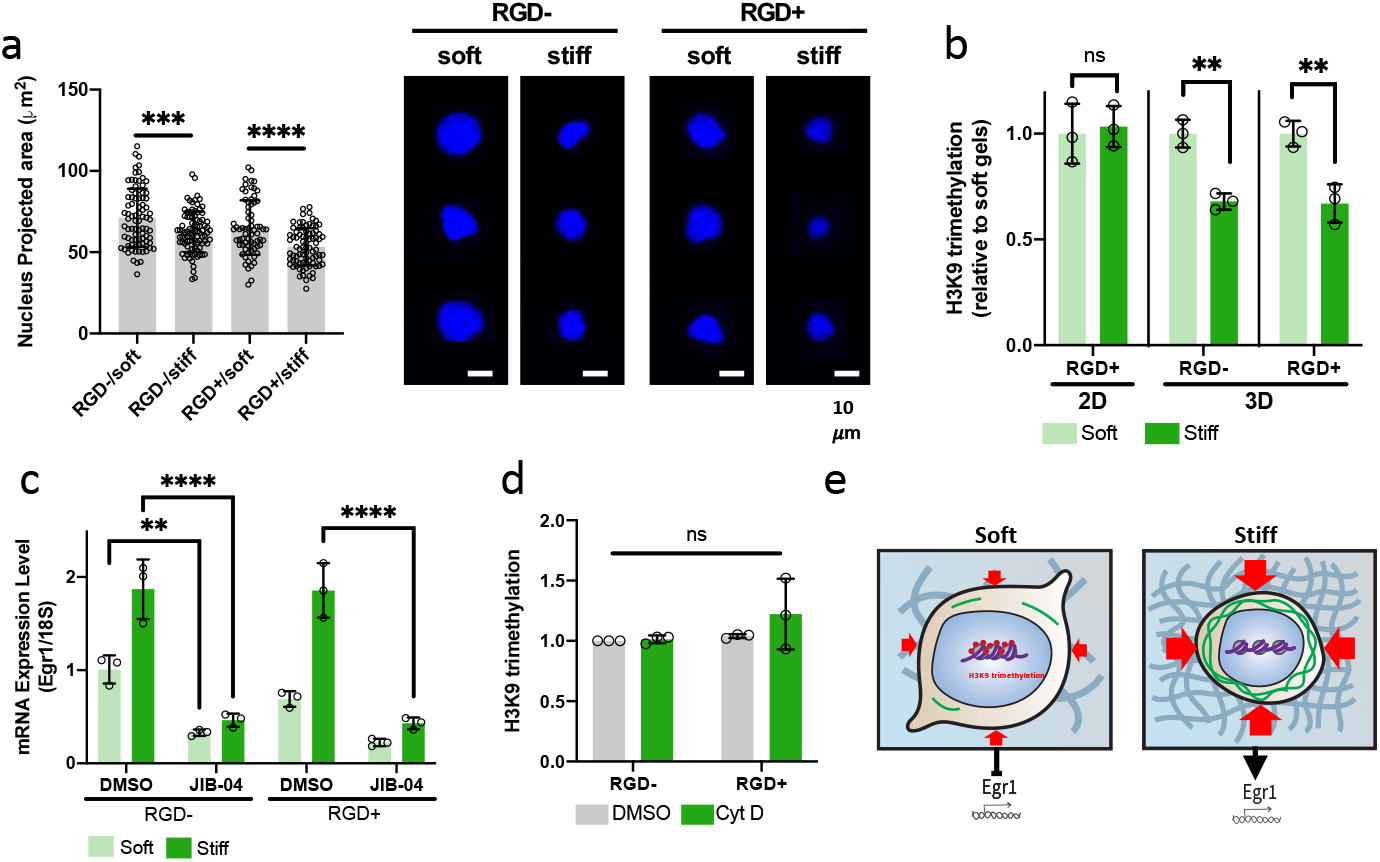
H3K9 trimethylation is associated with the stiffness-dependent *Egr1* expression in 3D matrices. **a**, Projected area of the cell nuclei in the four different 3D gels (RGD-/soft, RGD-/stiff, RGD+/soft, RGD+/stiff) (left) and the corresponding representative images of DAPI-stained cells in the gels after dissection. n>69 cells were used for each condition. Scale bar 10 *μ*m. **b**, Stiffness-dependence of H3K9me3 level in 2D and 3D gels (n=3). **c**, mRNA expression level of *Egr1* in the four different 3D gels after treatment of DMSO (control) and JIB-04 (3 *μ*M) for 3 hr. All the values are relative to the level under DMSO-treated RGD-/soft condition (n=3). **d**, H3K9me3 level of the cells in RGD- and RGD+ stiff gels after treatment of DMSO (control) and cytochalasin D (1 *μ*M). **e**, Schematics of the suggested mechanism of how *Egr1* expression is regulated by 3D gel stiffness-dependent confining stress. H3K9 trimethylation could be regulated by stiffness-dependent confining stress leading to the stiffness-dependent *Egr1* expression in 3D matrices. One-way ANOVA followed by Tukey test \*\*\*\**p*<0.001, \*\*\**p*<0.005, \*\**p*<0.01, \**p*<0.05. Graphs show mean ± s.d.

## DISCUSSION

We investigated whether, and by what mechanism, NSCs exhibit mechanosensitive differentiation in 3D. Using a crosslinked HA-DBCO gel, we observed that ECM stiffness regulates 3D mechanosensitive fate decisions in a narrower and more brain-mimetic stiffness range than 2D. Through unbiased transcriptome analysis, we identified a 3D matrix-specific mechanosensitive regulator, *Egr1*, whose genetic perturbation established its critical role in stiffness-dependent suppression of β-catenin signaling and neuronal differentiation. Furthermore, our results are consistent with the hypothesis that stiff 3D gels suppress neurogenesis due to enhanced confinement stress during volumetric expansion, which modulates actin assembly and increases *Egr1* expression.

While several studies have shown differences in mechanosensitive stem cell behaviors between 2D vs 3D microenvironments^5, 18, 19^, little is known about how dimensionality-specific physical factors and their underlying biomolecular mechanisms regulate stem cells. In this study, we identified *Egr1* as a key a mechanoresponsive transcriptional factor that regulates stiffness-dependent NSC lineage commitment. Intriguingly, *Egr1* exerted functional effects in 3D but not in 2D, where cells exhibited negligible *Egr1* expression. To our knowledge, *Egr1* thus represents the first reported 3D matrix-specific mechanosensitive stem cell regulatory factor. Higher expression of *Egr1* in 3D stiff gels suppressed the expression of *Prkaca* and in the activation of β-catenin signaling, raising the possibility of 3D-specific *Egr1* regulation of β-catenin signaling in numerous other biological processes.^51, 52^ For example, *Egr1* is known to regulate the synaptic plasticity and activity of mature neuronal circuits,^29, 31, 32^ and if it is also mechanosensitive in this context, it could modulate neuronal activity in older brain, which is known to soften with aging.

Another intriguing finding was that the stiffness-dependent *Egr1* expression was highly associated with a property specific to 3D matrix-specific mechanics (ECM confining stress), potentially explaining why *Egr1* functions only in 3D. Osmotic restriction of cell volumetric growth in stiff 3D gels, which would be expected to reduce confining stress, altered actin cytoarchitecture and attenuated *Egr1* expression. Thus, confining stress appears to increase *Egr1* expression and influence stiffness-dependent lineage commitment through a mechanism that involves altered actin cytoarchitecture.

Unexpectedly, the stiffness-dependent NSC transcriptome in general (**Fig. 2b**) and regulation of *Egr1* expression specifically was observed in both bare gels and gels functionalized with an RGD peptide and occurred even upon inhibiting protein secretion by Exo-1. Of course, these findings do not rule out the possibility that adhesion to RGD motifs, secreted matrix, or the HA backbone itself regulates other important biological functions besides lineage commitment ^53,54^. Instead, our model points to the centrality of confining stress in regulating lineage commitment, which is dictated by the mechanical properties of the surrounding matrix and not particular mechanism of adhesion.

Taken together, our work establishes an *Egr1*-mediated mechanotransduction pathway that controls lineage commitment through mechanisms that are intrinsic to 3D matrices. In the future, it would be fruitful to develop strategies to directly manipulate confining stress independently from the stiffness in 3D matrices to more precisely isolate the effect of confining stress on actin assembly, chromatin modification, and *Egr1* expression. It will also be important to more thoroughly identify the molecular mechanisms that link actin assembly, *Egr1* expression, β-catenin, and neurogenic lineage commitment.

## ACKNOWLEDGMENTS

We thank Olivia Teter (University of California, San Francisco) for experimental assistance. This work was supported by National Institutes of Health (R01NS074831 to S.K. and D.V.S.) and a Siebel Fellowship to J.B. RNA sequencing was carried by the DNA Technologies and Expression Analysis Core at the UC Davis Genome Center, supported by NIH Shared Instrumentation Grant 1S10OD010786-01.

## AUTHOR CONTRIBUTIONS

J.B. designed and performed experiments, analyzed and interpreted the data, and wrote the manuscript. P.A.L. contributed to RNA sequencing analysis and the experiment for generating *Egr1* knockdown cells. S.L. and T.-S.K. conducted ABAQUS simulation. S.K. and D.V.S. supervised the project, helped in experimental design and data interpretation, and co-wrote the manuscript.

## ADDITIONAL INFORMATION

**Supplementary information** is available for this paper.

**Reprints and permissions information** is available at www.nature.com/reprints.

**Correspondence and requests for materials** should be addressed to S.I.

### Publisher’s note

Springer Nature remains neutral with regard to jurisdictional claims in published maps and institutional affiliations.

