## Supplementary Information for "3D Microenvironment-Specific Mechanosensing Regulates Neural Stem Cell Lineage Commitment"

**List of Contents**

**1. Materials and Methods**
**2. Supplementary Figures S1–S12**
**3. Supplementary Table 1**

*Materials and Methods*

**Cell culture and differentiation.** Adult rat hippocampal NSCs were derived from adult female Fischer 344 rats (Charles River) as previously described[1]. The cells were cultured in Dulbecco's modified Eagle's Medium/nutrient mixture F-12 (DMEM-F12, Gibco) supplemented with N2 supplement (Life Technologies) and 20 ng/mL of FGF-basic (Peprotech) on the tissue-culture polystyrene plates coated with poly-ornithine (10 μg/mL, Sigma Aldrich) and laminin (5 μg/mL, Invitrogen) sequentially. The growth medium for undifferentiated cells was replenished every 2 days. To attain the cells in differentiated state, the cells were cultured in mixed differentiation medium (DMEM-F12 with N2 supplemented with 1 μM of retinoic acid and 1% FBS) right after being seeded onto gels or encapsulated with gels. The media was replaced every 2 days during 7 days of differentiation process. Then the cells were fixed for immunofluorescence imaging.

**DBCO functionalization to HA.** Before to make hydrogels, hyaluronic acid (HA) was modified with dibenzocyclooctyne (DBCO). First, carboxylic acid groups of sodium hyaluronate (average molecular weight 75 kDa, Lifecore Biomedical) dissolved in 2-(N-Morpholino)ethanesulfonic acid (MES) at 1 mg/mL were activated by N-(3-Dimethylaminopropyl)-N’-ethylcarbodiimide (EDC, Sigma Aldrich) and N-hydroxysuccinide (NHS, Sigma Aldrich) for 1 hr. Then, 0.6 equivalents of DBCO-amine (Sigma Aldrich) in dimethylsulfoxide (DMSO) were added dropwise to the solution. After 48 h reaction at room temperature, unreacted starting materials in the mixture were removed by centrifugation with a 10 kDa-cutoff concentrator (Millipore), and the remaining reaction mixture was precipitated and washed with cold acetone for twice. The precipitate was dissolved in ultrapure water and lyophilized for 2 days. The extent of DBCO functionalization to HA was estimated by 1H nuclear magnetic resonance (NMR) spectroscopy.

**Making HA-DBCO hydrogels.** DBCO-functionalized HA (HA-DBCO) formed hydrogel by strain-promoted azide alkyne cycloaddition (SPAAC), one of bio-orthogonal gelation. HA-DBCO was dissolved in DMEM/F-12 to a final concentration of 3 w/v%, and Poly(ethylene glycol) bisazide (PEG-bisazide, average $M_n$ 1,100, Sigma Aldrich) was added for crosslinking. RGD sequence-containing peptide with azide functionality (K($N_3$)GSGRGDSPG, 1mM, Genscript) was also added to the HA formulation at the same time as necessary. Then, the mixture was incubated for 10 min at 37 $^oC$ for crosslinking reaction. The amount of HA-DBCO and PEG-bisazide were varied to obtain the gels with different stiffnesses. For encapsulating the cells within hydrogels, cell suspension was added to the HA-DBCO solution right before adding PEG-bisazide.

**Hydrogel characterization.** Stiffnesses of the hydrogels were characterized by shear rheology via Physical MCR 301 rheometer (Anton Paar) with an 8-mm parallel plate geometry. Temperature of the gel solution was controlled with Peltier element (Anton Paar), and water droplet was added around solvent trap to prevent sample dehydration. Elastic moduli map was also characterized by atomic force microscopy (AFM) on a Vaeco (Bruker) Catalyst Bioscope instrument. The gel sample was soaked in cell culture medium (DMEM/F-12) at room temperature for all measurements. The deflection sensitivity of each MLCT-Bio cantilever was measured against a glass cover slide, and the spring constant was obtained by thermal tuning. Sneddon indentation model (cone indenting infinite half-space) was used to fit force-indentation curves for calculating elastic moduli.

**Immunocytochemistry.** Samples were fixed by incubating with 4 % paraformaldehyde in PBS for 15 min at room temperature. After thorough washing with PBS, the fixed cells were permeabilized and blocked with Triton X-100 (0.3 %) and BSA (3 %) in PBS for 35 min at RT. Then the samples were incubated at 4 $^{o}$C for 48 hr with the following primary antibodies: mouse anti-Tubulin β3 (TUBB3) (1:1000, BioLegend), rabbit anti-glial fibrillary acidic protein (GFAP) (1:1000, Abcam), rabbit anti-early growth response 1 (Egr1) (1:1000, Cell Signaling Technology), rabbit anti-β-catenin (Cell Signaling Technology). After washing with PBS, the resulting samples were stained with goat anti-rabbit IgG (H+L) secondary antibody or Alexa Fluor 546 (Invitrogen), goat anti-mouse IgG (H+L) secondary antibody, Alexa Fluor 633 conjugate (Invitrogen) for 40 min at RT. Nuclei were labeled with 4′,6-diamidino-2-phenylindole (DAPI, Sigma Aldrich) and F-actin was labeled with Alexa Fluor 546 Phalloidin (Thermo-Fisher) if necessary. All fluorescent images were taken using Prairie Technologies 2-photon and confocal microsope, QuantEM 512SC camera, 60X and 40X objective lenses, and native Prairie View software and visualized by z-stack mode. All the image analysis and processing was performed with ImageJ and Imaris.

**Sectioning Samples.** After fixation, the hydrogels containing cells were embedded in OCT-compound solution (Tissue-Tek) at -80 $^{o}$C for 30 min and sectioned to the thickness of 15-30 μm via a cryostat NX50 (ThermoFisher). Sectioned samples were used to characterizing β-catenin nuc/cyt ratio and measuring actin intensity after immunostaining.

**Blocking cell-ECM interactions.** For blocking RGD-binding integrins, NSCs were preincubated for 30 min with 500 μg of RGD sequence-containing peptide (GRGDSP, Bachem) and control (GRADSP, Bachem) before encapsulation with hydrogel. After encapsulation, mixed differentiation medium containing the peptides were added to each well. Optimal concentration of the peptides was determined by cell adhesion assay. The peptides-treated cells with varied peptide concentrations were seeded onto the RGD peptide-ligated (1 mM) HA-DBCO hydrogels (1 kPa). After 24 hr, the samples were washed twice with PBS, and the number of adhered cells was counted for each concentration condition. For inhibiting the protein secretion of cells, 150 μM of 2-(4Fluorobenzoylamino)-benzoic acid methyl ester (Exo-1, Sigma Aldrich) dissolved in DMSO was treated to the cells in mixed differentiation medium right after encapsulation. DMSO treatment was performed for control. Optimal concentration of Exo-1 was obtained by Live/dead assay with varied Exo-1 concentrations after 24 hr.

**RNA isolation and qPCR.** NSCs were extracted from HA-DBCO gels by incubating the gels in DMEM/F-12 containing hyaluronidase (750-3000 U/mL, Sigma Aldrich) for 30 min at 37 $^{o}$C. The suspensions were centrifuged at 200×g for 2 min to pellet cells and washed with PBS. Phenol-free total RNA was extracted from the cell pellets using RNeasy Plus Micro Kit with gDNA eliminator columns (Quiagen) following the manufacturer's protocol. After measurement of total RNA concentration, 600 ng of RNA was converted to cDNA using an iScript cDNA synthesis kit (Bio-Rad). Obtained cDNA was used for SYBR Green (Bimake) quantitative polymerase chain reaction (qPCR) with a 5 μM final forward and reverse primer concentration. Primer sequence was listed in **Supplementary table 1**. qPCR was conducted for 34 cycles in a CFX connect real-time PCR system (Bio-Rad). RNA level analysis was performed by $ddC_t$ method, and each gene expression was internally normalized by the expression level of housekeeping gene (S18)[2] run on the same qPCR batch.

**RNA sequencing and data analysis.** RNA was isolated and purified as described above with two biological replicates per each hydrogel condition (RGD-/soft, RGD-/stiff, RGD+/soft, RGD+/stiff). RNA integrity number (RIN) was assessed by using Agilent 2100 Bioanalyzer (Agilent Technologies), and the high quality of RNA (RIN $\geq$ 9.8) was used. RNA-seq library preparation and batch-tag-RNA-seq (3'-tag-seq gene expression profiling) were carried out by the DNA Technologies and Expression Analysis Core at the University of California, Davis Genome Center. The reads were initially trimmed using BBDuk to remove Illumina adapters and to filter low quality reads. The samples were then aligned to ratRibosomal genes using bbmap to remove ribosomal contaminated RNA. Any samples less than 21 bases were removed using BBduk. Read quality analysis was performed using FastQC to assure there was a Phred score > 28. The reads were aligned to the rattus norvegicus genome (rn6) using hisat2 v2.1.0. Then the aligned reads were converted to counts using htseq. Normalization and differential expression analysis were conducted using DESeq2 and other Bioconductor packages in R. Differentially expressed genes (DEGs) were determined with the condition of *P*-value less than 0.05. Heatmaps and volcano plots were also obtained by Bioconductor packages in R.

**Western blotting.** NSCs embedded in HA-DBCO hydrogels were isolated by treating hyaluronidase (750-3000 U/mL, Sigma Aldrich) at 37 $^{o}$C for 30 min as previously described above. Total protein was extracted from the cell pellet washed with PBS once by lysing with RIPA lysis buffer (Sigma Aldrich) containing Halt$^{TM}$ proteinase and phosphatase inhibitor cocktail (Thermo Scientific) on ice for 10 min. The protein concentration was determined by Bicinchoninic acid (BCA) assay with Pierce$^{TM}$ BCA protein assay kit (Thermo Scientific), and samples were normalized with respect to protein content. Proteins were separated via SDS-PAGE and transferred onto a nitrocellulose membranes (0.22 μm, Odyssey). Membranes were blocked in a tris-buffered saline (TBS) blocking buffer (Odyssey) for 40 min and incubated with primary antibodies [rabbit anti-Egr1 (1:1000, Cell Signaling Technology), mouse anti-Yes-associated protein (YAP) (1:1000, Cell Signaling Technology), rabbit anti-Axin1 (1:1000, Invitrogen), mouse anti-β-actin (1:10000, Sigma Aldrich)] overnight at 4 $^{o}$C. Then, the membranes were washed with TBST and treated with biotinylated secondary antibodies [goat anti-rabbit IgG H&L (biotin) (1:10000, Abcam) or goat anti-mouse IgG H&L (biotin) (1:10000, Abcam)] and streptavidin-conjugated fluorescent label [Streptavidin, Alexa Fluor 790 conjugate (Invitrogen) or Streptavidin, Alexa Fluor 700 conjugate (Invitrogen)], sequentially. The membranes were washed with TBST and imaged by Odyssey CLx (LI-COR Biosciences).

**shRNA cloning.** shRNA inserts were designed with AgeI- and EcoRI-based overhangs using Ensembl genome browser and the online tool, InvivoGen. shRNA targeting rat Egr1 (shEgr1-1, GCCGAGATGCAATTGATGTCT; shEgr1-2, GTCGAATCTGCATGCGTAATT) and a scramble control (GTCGGCTACGAAGGTATTCTA) were obtained from Elim Biopharmaceuticals. The inserts were ligated into the pLKO.1 puro vector (Addgene plasmid #10878).

**Viral packaging and transduction.** Lentiviral particles were packaged in HEK 293T cells with psPAX2 and pMD2.G through polyethylenimine (PEI) transfection and purified as previously described[3]. Purified viral particles were transduced to the NSCs with a multiplicity of infection (MOI) of 1, and shRNA-expressing cells were selected using 1 μg/mL puromycin longer than 4 days.

**Nuclear to cytoplasmic ratio quantification.** Images from 30-μm sectioned samples co-stained for total β-catenin, nuclei and F-actin were used to quantify β-catenin nuclear-to-cytosolic ratio based on fluorescence intensity. Binary masks of the nuclei and actin were obtained from DAPI and Phalloidin images and superimposed each other to create masks that contain the cytosol but exclude the nucleus. Then the total β-catenin fluorescence intensity was quantified in these regions and their ratio was calculated followed by normalizing to the area of each domain.

$$\beta-\text{catenin}\ \frac{\text{nuc}}{\text{cyt}}\ ratio = \frac{\frac{integrated\ intensity\ of\ nuclear\ \beta-\text{catenin}}{area\ of\ nucleus}}{\frac{integrated\ intensity\ of\ cytosolic\ \beta-\text{catenin}}{area\ of\ cytosol}}$$

**Luciferase assay.** Naïve NSCs and Egr1 knock down (KD) NSC cell lines were transduced with a lentiviral construct encoding a 7×TFP TFC/LEF luciferase reporter, which represents β-catenin-TCF/LEF-based transcription. Cell pellets from the cells encapsulated with HA-DBCO gel under each condition were obtained after 3 days of encapsulation/differentiation as described above. The cell pellets were washed with PBS once, and lysed with lysis buffer (Promega), and centrifuged to pellet debris. Suspensions were loaded to each well of a white opaque 96-well plate, and Luciferase Assay Reagent (Promega) was treated right before detection. Luminescence intensity was detected via SpectraMax luminometer (Molecular Devices), and normalized to total protein concentration obtained through BCA protein assay (Pierce) to consider the variation of proliferation between samples.

**Image Analysis.** Actin intensity was quantified by extraction of intensity line-scans perpendicular to the membrane by linearizing of the cell edges using Fiji. Rhodamine phalloidin-stained fluorescent images for the cells encapsulated within 3D gels for 5 hr after gel dissection were used for the analysis. Obtained intensity line-scans were fitted and then the peak intensity was measured through Origin data analysis software to quantify actin intensity. Full width at half maximum (FWHM) of the Gaussian whose peak was located closest to the center of the line was used as a measure of thickness by Origin. Cellular volume was measured by using Imaris 3D imaging software. Fluorescent images showing 3D rendering of single NSCs stained with cell membrane dye (R18) were used to quantify the volume.

**Abaqus simulation.** In order to analyze the stress distribution of the hydrogel with the growth of the cell, the finite element analysis (FEA) was conducted using a nonlinear static solver in ABAQUS 6.14. The sphere-shaped cell and the hexahedron-shaped hydrogel were modeled as a 3D deformable solid, and the cell was designed to be located at the center of the hydrogel. The experimentally obtained cellular volume was used as cell sizes, and the length of each side of the hydrogel was assumed as 100 μm. The element type of the cell and the hydrogel was a C3D20R (A 20-node quadratic brick, reduced integration). The elastic modulus of the hydrogel, assumed to be isotropic, was obtained with its shear modulus and Poisson's ratio: $E=2G(1+\nu)$. For the calculation, the shear modulus values of each hydrogel obtained by rheometer were used, and the Poisson's ratio was assumed to be 0.49. Meanwhile, the elastic modulus and Poisson's ratio of the cell were designated to have constant values of 500 Pa and 0.49, respectively.

**Inhibition experiments.** In order to inhibit myosin II, FAK, and actin assembly, NSCs were preincubated with blebbistatin (1 μM, Sigma Aldrich), PF-573228 (0.5 μM, Sigma Aldrich), and cytochalasin D (1 μM, Sigma Aldrich), respectively, for 20 min. Right after gelation, these were added to the media with same concentration under the spontaneous differentiation condition. Then the cells were harvested after 5 hr to investigate whether each inhibition affect Egr1 mRNA expression. JIB-04 (3 μM, Cayman Chemical Company), pan-selective Jumonji histone demethylase inhibitor, was treated to cell media right after encapsulation of NSCs with 3D gels to see the effect of H3K9me3 on *Egr1* expression.

**H3K9me3 quantification.** Histone lysates were isolated from the cells ($1x10^6$) incubated within each hydrogel for 3 hr under differentiation condition by Epiquik$^{TM}$ Total Histone Extraction Kit (Epigentek), and the quantification of H3K9me3 level was performed with EpiQuik Global Tri-Methyl Histone H3K9 Quantification Kit (Epigentek) following the protocols provided by manufacturer.

**Statistical analysis.** All the quantitative data were presented as the mean ± standard deviation (s.d.) and the number of biological and technical replicates are indicated in the figure legends and methods section. One-way ANOVA followed by Tukey test and Student's *t*-test for between-group differences were performed with GraphPad Prism as indicated in the figure legends.

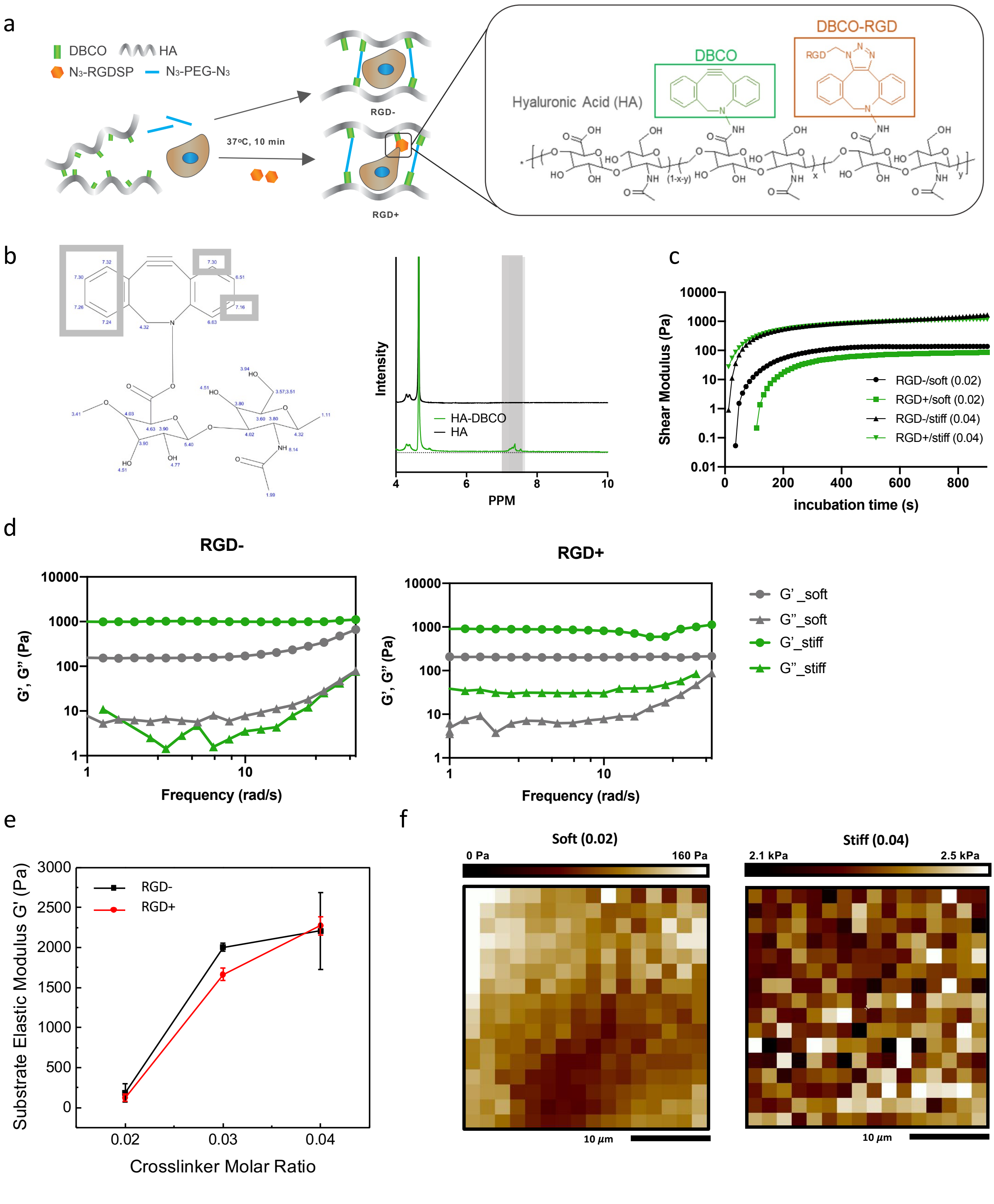


**Supplementary Figure S1.** **a**, Schematic of the encapsulation of NSCs using HA-DBCO hydrogels with and without RGD functionalization. **b**, $H^1$ Nuclear magnetic resonance (NMR) spectroscopy of bare HA and DBCO-functionalized HA. **c**, Shear elastic modulus change of HA-DBCO hydrogels with incubation time. **d**, Frequency sweep of the hydrogels: RGD-/soft and RGD-/stiff (left); RGD+/soft and RGD+/stiff (right). **e**, Substrate elastic moduli of RGD- and RGD+ HA-DBCO hydrogels with the ratio of azide to HA monomers (0.02-0.04) measured by atomic force microscopy (AFM) nanoindentation. n=3. **f**, The modulus distribution map of RGD-/soft (left) and RGD-/stiff (right) HA-DBCO gels characterized by AFM.

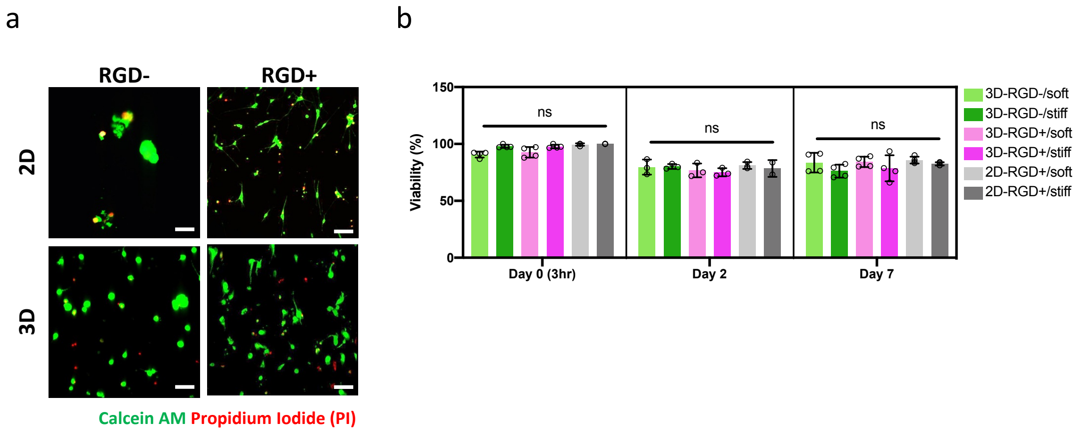


**Supplementary Figure S2. a**, Cell viability assay of the NSCs on 2D and in 3D gels with and without RGD functionalization under spontaneous differentiation condition for 2 days. The cells stained by Calcein AM (green) and propidium iodide (PI) (red) are live and dead cells, respectively Scale bar, 50 µm. b, Quantification of the number percentage of live cells after differentiation for 0 hr, 2 days, and 7 days. n= 3 biological replicates were used. One-way ANOVA followed by Tukey test *$p$<0.05. Graphs show mean ± s.d.

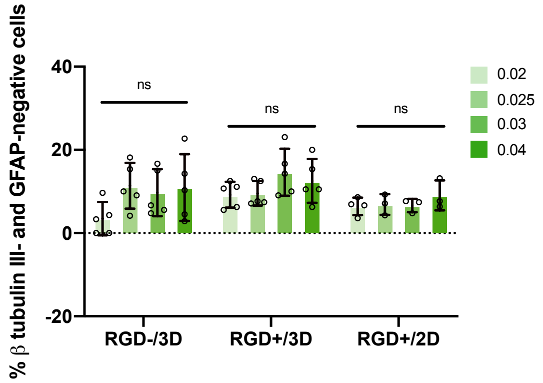


**Supplementary Figure S3.** Number percentage of the cells negative for both β-tubulin III and GFAP encapsulated within RGD- and RGD+ 3D gels and RGD+ 2D gels with the different molar ratio of azide to HA monomer: 0.02, 0.025, 0.03, and 0.04. n=3 to 5 biological replicates were used. One-way ANOVA followed by Tukey test **$p$<0.01, *$p$<0.05. Graphs show mean ± s.d.

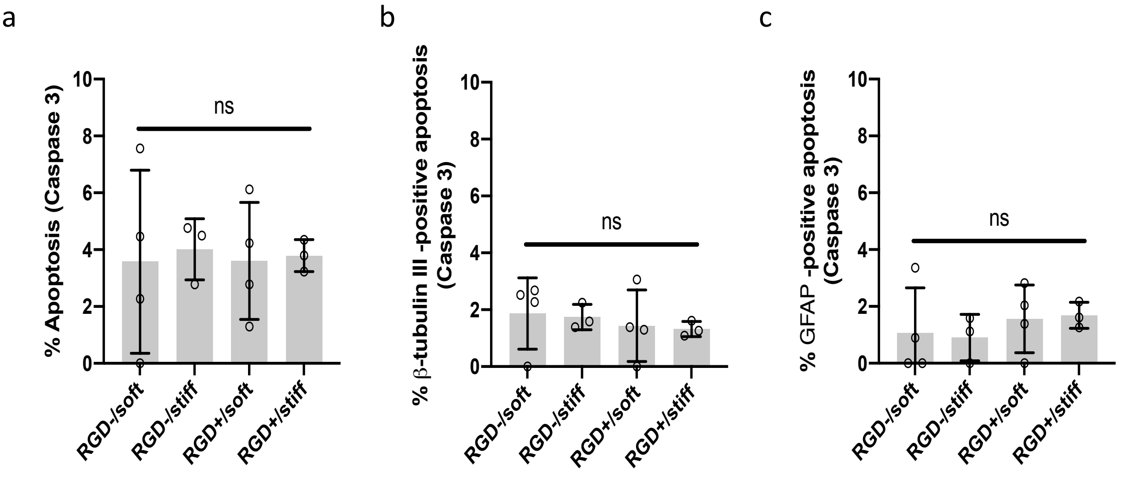


**Supplementary Figure S4.** Number percentage of the NSCs (**a**), β-tubulin III+ NSCs (**b**), GFAP+ NSCs (**c**) positive for Caspase 3 (apoptotic) in four different 3D HA-DBCO gels: RGD-/soft, RGD-/stiff, RGD+/soft, RGD+/stiff. The cells were differentiated for 7 days. n=3 to 4 biological replicates were used. One-way ANOVA followed by Tukey test **p*<0.05. Graphs show mean ± s.d.

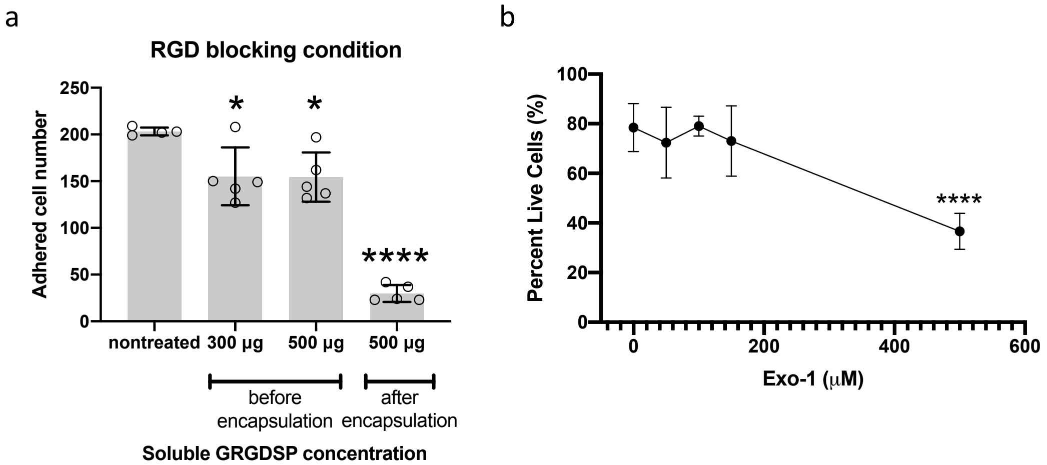


**Supplementary Figure S5. a**, Quantification of the number of NSCs adhered on RGD-functionalized substrates after treatment of RGD sequence-containing peptide for blocking RGD-integrin binding. The most right represents the condition that 500 μg of RGD peptide was pretreated before encapsulation and added to the media for 24 hr after encapsulation as well n=5 replicates were used with two biological replicates. **b**, Cell viability assay after treatment of Exo-1 with various concentrations. One-way ANOVA followed by Tukey test *****p*<0.001, **p*<0.05. Graphs show mean ± s.d.

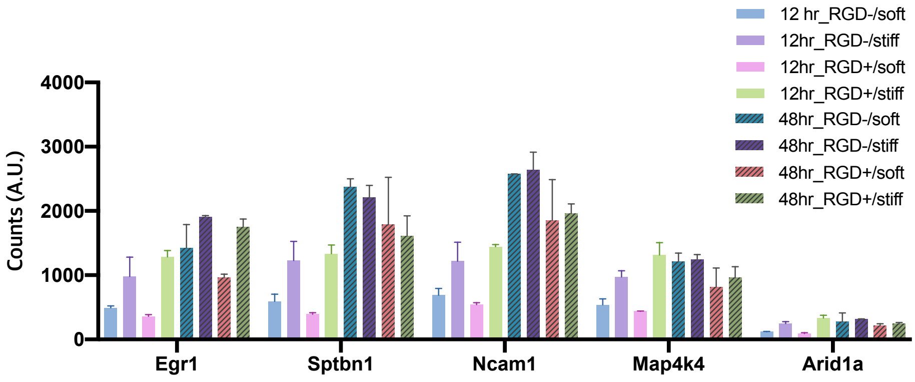


**Supplementary Figure S6.** Transcript counts obtained from RNA sequencing analysis for the top five DEGs (Egr1, Sptbn1, Ncam1, Map4k4, Arid1a) from stiff vs. soft gel comparisons. RNA was extracted from NSCs differentiated for 12 hr and 48 hr in the four different hydrogels: RGD-/soft, RGD-/stiff, RGD+/soft, RGD+/stiff. All the five DEGs exhibited strong stiffness-dependence only after 12 hr of differentiation in the gels.

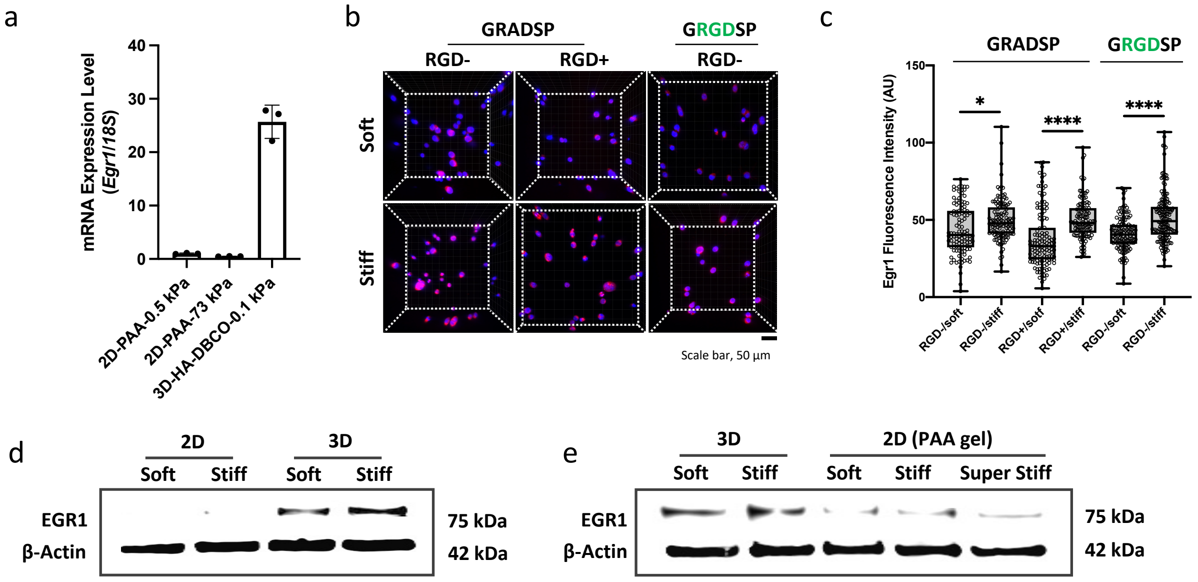


**Supplementary Figure S7. a**, *Egr1* mRNA expression of NSCs in 2D polyacrylamide (PAA) gels and 3D HA-DBCO gels. Representative immunofluorescent staining of EGR1 (**b**) and quantification of EGR1 fluorescence intensity (**c**) for the NSCs differentiated in RGD- soft and stiff hydrogels with the treatment of RAD- (control) and RGD- containing soluble peptides (500 µg) for 24 hr. Scale bar 50 µm. n>108 cells per each condition was analyzed for the quantification of EGR1 fluorescence intensity. Western blotting of EGR1 in the NSCs differentiated in 2D and 3D HA-DBCO hydrogels (**d**) and in 3D HA-DBCO gels and 2D PAA gels with higher stiffness (for 2D, superstiff – 73 kPa) (**e**) for 24 hr. Soft: 0.1 kpa, Stiff: 1.2 kPa. One-way ANOVA followed by Tukey test ****$p$<0.001, *$p$<0.05. Graphs show mean ± s.d.

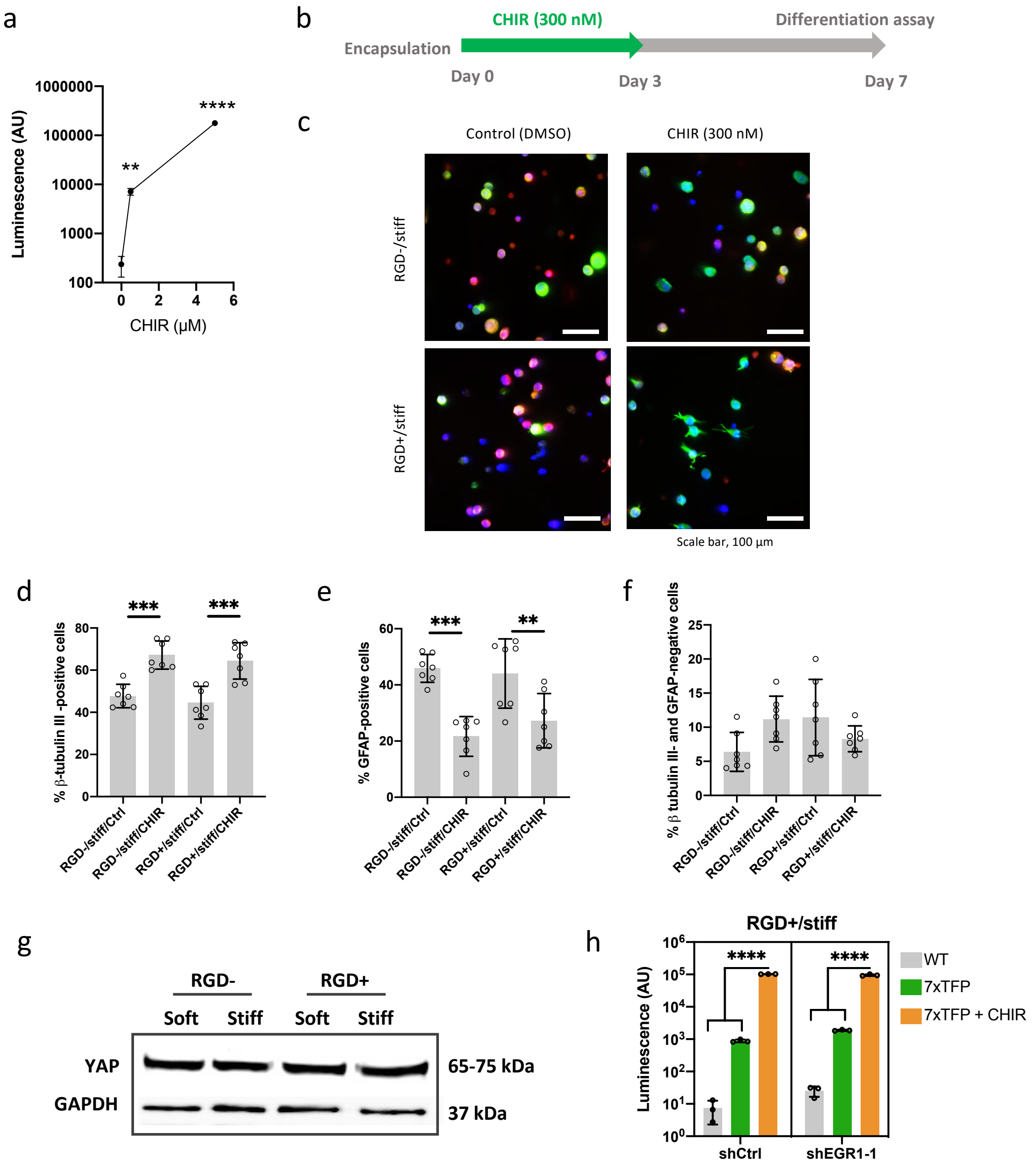


**Supplementary Figure S8. a**, Luciferase assay for the NSC cell line carrying 7xTFP reporter for β-catenin activity treated with DMSO (control) and CHIR (0.5 μM, 5 μM) for 3 days. **b**, Experimental timeline of CHIR (300 nM) treatment for differentiation assay to investigate if β-catenin signaling plays a role in NSC lineage commitment in 3D gels. **c**, Representative images of immunostaining for β-tubulin III (green), GFAP (red), DAPI (blue) in the RGD-/stiff and RGD+/stiff gels. The NSCs were treated with DMSO (control) or CHIR (300 nM) as shown in **b**. Scale bar 100 μm. Quantification of β-tubulin III- (**d**) and GFAP- (**e**) positive cells and the cells negative for both β-tubulin III and GFAP (**f**). n=5 with three biological replicates were used. **g**, Western blotting of total YAP in the NSCs differentiated in four different 3D gels (RGD-/soft, RGD-/stiff, RGD+/soft, RGD+/stiff) for 24 hr. **h**, Luciferase assay for shCtrl and shEGR1-1 NSC cell lines carrying 7xTFP reporter for β-catenin activity. The cells were treated with DMSO (control) or CHIR (0.5 μM) in RGD+/stiff 3D gels for 3 days under spontaneous differentiation condition. Wild type (WT) NSCs that do not carry the 7xTFP were used as negative control. One-way ANOVA followed by Tukey test ****$p$<0.001, ***$p$<0.005, **$p$<0.01. Graphs show mean ± s.d.

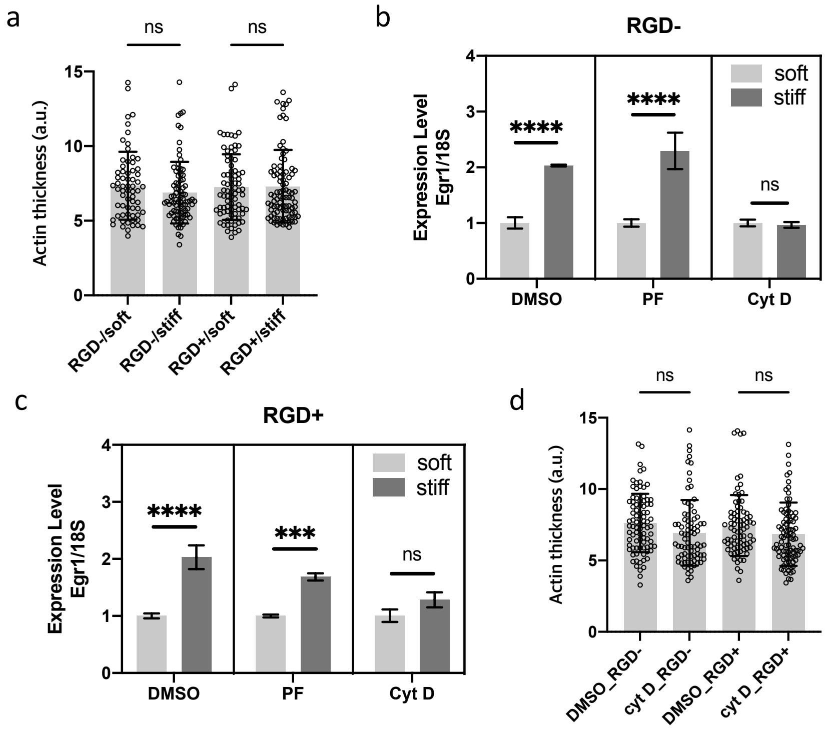


**Supplementary Figure S9. a**, Quantification of relative peak cortex actin thickness of the NSCs encapsulated within 3D gels for 5 hr. Stiffness-dependence of *Egr1* mRNA expression in RGD- (**b**) and RGD+ (**c**) 3D gels after treatment of DMSO, PF-573228 (0.5 μM), and cytochalasin D (1 μM) for 5 hr. The expression level is relative to that on soft gel for each condition. n=3. **d**, quantification of relative peak cortex actin thickness after treatment of DMSO (control) and cytochalasin D (1 μM) in RGD-/stiff and RGD+/stiff gels for 5 hr. 66-87 cells were used for the quantification. One-way ANOVA followed by Tukey test ****$p$<0.001, ***$p$<0.005. Graphs show mean ± s.d.

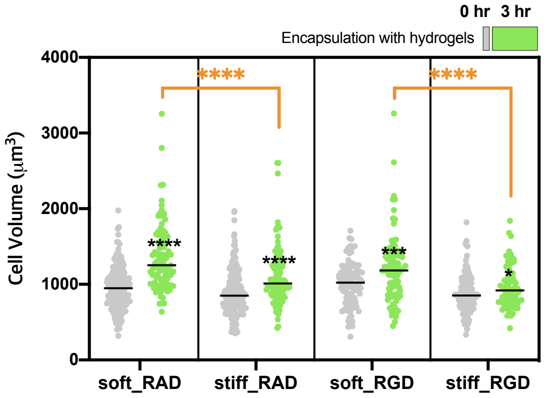


**Supplementary Figure S10. a**, Quantification of cell volumes after 0 hr and 3 hr of encapsulation in RGD-/soft and RGD-/stiff 3D gels after treatment of RAD- and RGD-containing peptides to investigate whether blocking RGD-integrin interaction influences cellular volume growth in 3D gels. This was measured from 3D rendering of single NSCs stained with cell membrane dye (R18), and the cells were embedded in under differentiation condition right after the encapsulation. 77-170 cells were used for the quantification of each condition. Scale bar 30 μm. One-way ANOVA followed by Tukey test ****$p$<0.001, ***$p$<0.005, *$p$<0.05. Graphs show mean ± s.d.

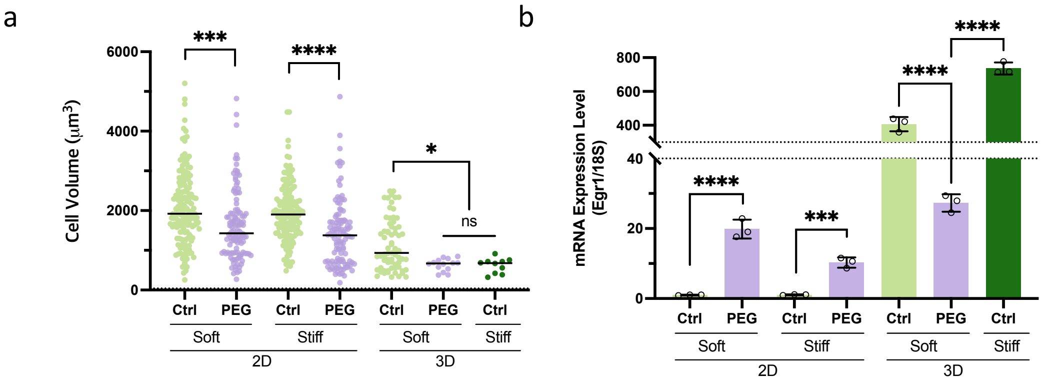


**Supplementary Figure S11.** Quantification of cell volumes (**a**) and mRNA expression level of *Egr1* (**b**) after 3 hr of encapsulation for the 2D and 3D gels under non-treated (Ctrl) and PEG 400 (1.5 wt%)-treated conditions. One-way ANOVA followed by Tukey test for each 2D and 3D gel groups ****$p$<0.001, ***$p$<0.005, *$p$<0.05. Graphs show mean ± s.d.

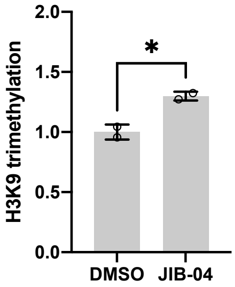


**Supplementary Figure S12. a**, H3K9me3 level of NSCs after treatment of DMSO and JIB-04 (3 μM) for 5 hr. One-way ANOVA followed by Tukey test *$p$<0.05. Graphs show mean ± s.d.

| Target | Primers |
|---|---|
| *Egr1* | F: GTATGCTTGCCCTGTTGAGTCC |
| | R: CATGCAGATTCGACACTGGAAG |
| *Axin1* | F: GACTTGGCCTTTGGTGGTAA |
| | R: AGGACGTTCAATGGACAAGG |
| *Prkaca* | F: GACCTCATCTACCGGGACCT |
| | R: TGTTGCCCCTTCCCCAGTAT |
| *Dvl1* | F: CCTACAAATTCTTCTTCAAGTCTATGG |
| | R: CTTGGCATTGTCGTCGAA |
| *18S* | F: GTAACCCGTTGAACCCCATTC |
| | R: CCATCCAATCGGTAGTAGCGA |

**Supplementary Table 1.** Primers used in qRT-PCR
